# Single-molecule imaging of telomerase RNA reveals a Recruitment – Retention model for telomere elongation

**DOI:** 10.1101/2020.01.31.929026

**Authors:** Hadrien Laprade, Emmanuelle Querido, Michael J. Smith, David Guérit, Hannah Crimmins, Dimitri Conomos, Emilie Pourret, Pascal Chartrand, Agnel Sfeir

## Abstract

Extension of telomeres is a critical step in the immortalization of cancer cells. This complex reaction requires proper spatio-temporal coordination of telomerase and telomeres, and remains poorly understood at the cellular level. To understand how cancer cells execute this process, we combined CRISPR genome editing and MS2 RNA-tagging to image single-molecules of telomerase RNA (hTR). Real-time dynamics and photoactivation experiments of hTR in Cajal bodies (CBs) reveal that hTERT controls the exit of hTR from CBs. Single-molecule tracking of hTR at telomeres shows that TPP1-mediated recruitment results in short telomere-telomerase scanning interactions, then base-pairing between hTR and telomere ssDNA promotes long interactions required for stable telomerase retention. Interestingly, POT1 OB-fold mutations that result in abnormally long telomeres in cancers act by enhancing this retention step. In summary, single-molecule imaging unveils the life-cycle of telomerase RNA and provides a framework to understand how cancer-associated mutations mechanistically drive defects in telomere homeostasis.

## Introduction

Mammalian telomeres consist of tracts of duplex TTAGGG repeats bound by a six-subunit protein complex – termed shelterin – that prevents the activation of the DNA damage response (de Lange, 2005; Sfeir and de Lange, 2012). Telomere repeats are synthesized by telomerase, a specialized ribonucleoprotein (RNP) that is minimally composed of a reverse transcriptase (hTERT) and an RNA subunit (hTR) (Greider and Blackburn, 1985). Most somatic cells lack telomerase activity and as a result, telomeres get progressively shorter until the cells become senescent or die. Reactivation of telomerase is a key event during tumorigenesis and enables cancer cells to proliferate indefinitely (Kim et al., 1994). Generating an active telomerase RNP is a multistep process that requires hTERT, hTR, and additional co-factors including dyskerin, TCAB1, and H/ACA proteins (Schmidt and Cech, 2015). Telomerase folding and assembly involves transit through Cajal bodies (CBs) (Jady et al., 2004; Tomlinson et al., 2006; Zhu et al., 2004) where processing and maturation of small nuclear RNAs (snRNAs) and snoRNPs take place (Gall, 2000). Trafficking of hTR through CBs relies on its interaction with the WD40 repeat-containing protein TCAB1 (Venteicher et al., 2009); (Cristofari et al., 2007) and is essential for proper targeting of telomerase to its substrate (Stern et al., 2012). Paradoxically, elimination of human coilin did not alter telomere length (Chen et al., 2015; Vogan et al., 2016), potentially implicating a backup pathway during telomerase assembly. It is also possible that other components of CBs compensate for coilin loss to allow the maturation and trafficking of hTR.

Following its exit from CBs, telomerase is targeted to chromosome ends to promote telomere elongation. It has been estimated that cancer cells have ∼250 active telomerase RNPs (Cohen et al., 2007) that add ∼50 nucleotides to most chromosome ends each cell cycle (Zhao et al., 2009). It is not fully understood how telomerase molecules are targeted to telomeres in a crowded nucleus. Emerging evidences indicate that telomerase recruitment is dependent on a specific interaction between hTERT and the shelterin subunit, TPP1. Disruption of TPP1-hTERT interaction leads to progressive telomere shortening and results in cell death (Nandakumar et al., 2012; Schmidt et al., 2014; Sexton et al., 2014; Zhong et al., 2012). In addition, single-particle tracking of hTERT-K78E, a mutant allele that is unable to interact with TPP1, confirmed that telomerase association with telomeres is diminished when hTERT-TPP1 interaction is blocked (Schmidt et al., 2016).

Telomere length is maintained around a set point that is essential for survival. Short telomeres trigger a DNA damage response that can lead to age-related diseases and short-telomere syndromes (also known as telomeropathies) (Martinez and Blasco, 2017). On the other hand, abnormally long telomeres can foster tumorigenesis. Telomere length regulation is not fully understood, but genetic evidence points to a role for DNA damage factors and telomere binding proteins in this process. For example, the ATM-like kinase, Tel1 and the ATR-like kinase, Mec1/Rad3 play partially redundant roles in maintaining telomere length in budding and fission yeasts (Moser et al., 2011; Ritchie et al., 1999). Inhibition of the ATM kinase in human and mouse cells compromises telomere repeat addition (Lee et al., 2015), and both ATM and ATR were reported to regulate telomerase localization at chromosome ends (Tong et al., 2015). Subunits of the shelterin complex have also been implicated in telomere length regulation (Loayza and De Lange, 2003; van Steensel and de Lange, 1997). Expression of POT1-ΔOB that is incapable of binding to single-stranded telomere DNA results in rapid telomere elongation (Loayza and De Lange, 2003). Notably, POT1 mutations have been reported in familial and sporadic cancers, and manifests abnormally long telomeres (Sfeir and Denchi, 2016). The mechanism by which POT1 mutations, which predominantly cluster in the DNA binding domain, alter repeat addition remains unknown.

Here, we apply the MS2-tagging approach to visualize single-hTR particles using live-cell imaging and photoconversion experiments. Our results fully delineate the dynamic process of telomerase trafficking through Cajal bodies. Furthermore, we provide evidence in support of a two-step “Recruitment – Retention” model that governs telomerase association with telomere ends. Telomerase recruitment by TPP1 gives rise to quick and highly diffusive associations. A subsequent retention step stabilizes telomerase at the 3’ overhang and promotes longer interactions with constrained diffusion. Interestingly, we show that POT1 OB-fold mutations enhances telomerase retention, thereby providing insight into the mechanism by which recently identified cancer-associated POT1 mutations lead to abnormal telomere elongation

## RESULTS

### MS2-tagging reveals hTR dynamics in live cells

To monitor the dynamics of human telomerase at the single molecule level, we employed the MS2-GFP tagging system and visualized its RNA subunit *in vivo*. The two-component MS2 system included a GFP-tagged MS2 coat protein (GFP-MCP) that recognizes the MS2 RNA stem-loop, even when added to a particular RNA (Bertrand et al., 1998; Querido and Chartrand, 2008). This system has been successfully employed to study the trafficking of RNA in many organisms and was used to visualize telomerase RNA in living yeast cells (Bertrand et al., 1998; Gallardo et al., 2011). Based on the recently reported Cryo-EM structure of telomerase holoenzyme, we anticipated that the 5’-end and the conserved regions 4 and 5 (CR4/5) domain of hTR could be modified with minimal disruption to the telomerase holoenzyme (Nguyen et al., 2018) (Figures 1A and S1A). To determine the number of MS2 stem-loop repeats that could be tolerated without compromising the function of the RNP, we employed a mutant hTR – termed hTR-TSQ1 – designed to add GTTGCG variant repeats that are distinguishable from the canonical telomere repeats (Diolaiti et al., 2013). We overexpressed hTR-TSQ1 tagged with MS2 stem-loops at the 5’-end and the CR4/5 region, independently, in 293T cells. Southern blot analysis was performed to detect incorporated GTTGCG sequences. Our results demonstrated that modification of hTR with > 4MS2 stem loops severely compromised repeat addition, whereas 3MS2 stem-loops did not impair the function of the RNP (Figure S1B-C). To further confirm these results, we expressed MS2-tagged hTR in VA13 cells that lack endogenous telomerase RNA, and monitored telomerase activity using Telomerase Repeated Amplification Protocol (TRAP) assay. While tagging hTR at the 5’end did not impair the function of the enzyme, the presence of 3MS2 stem loops at the CR4/5 domain slightly reduced telomerase activity *in vivo* (Figure S1D-G).

**Figure 1.**
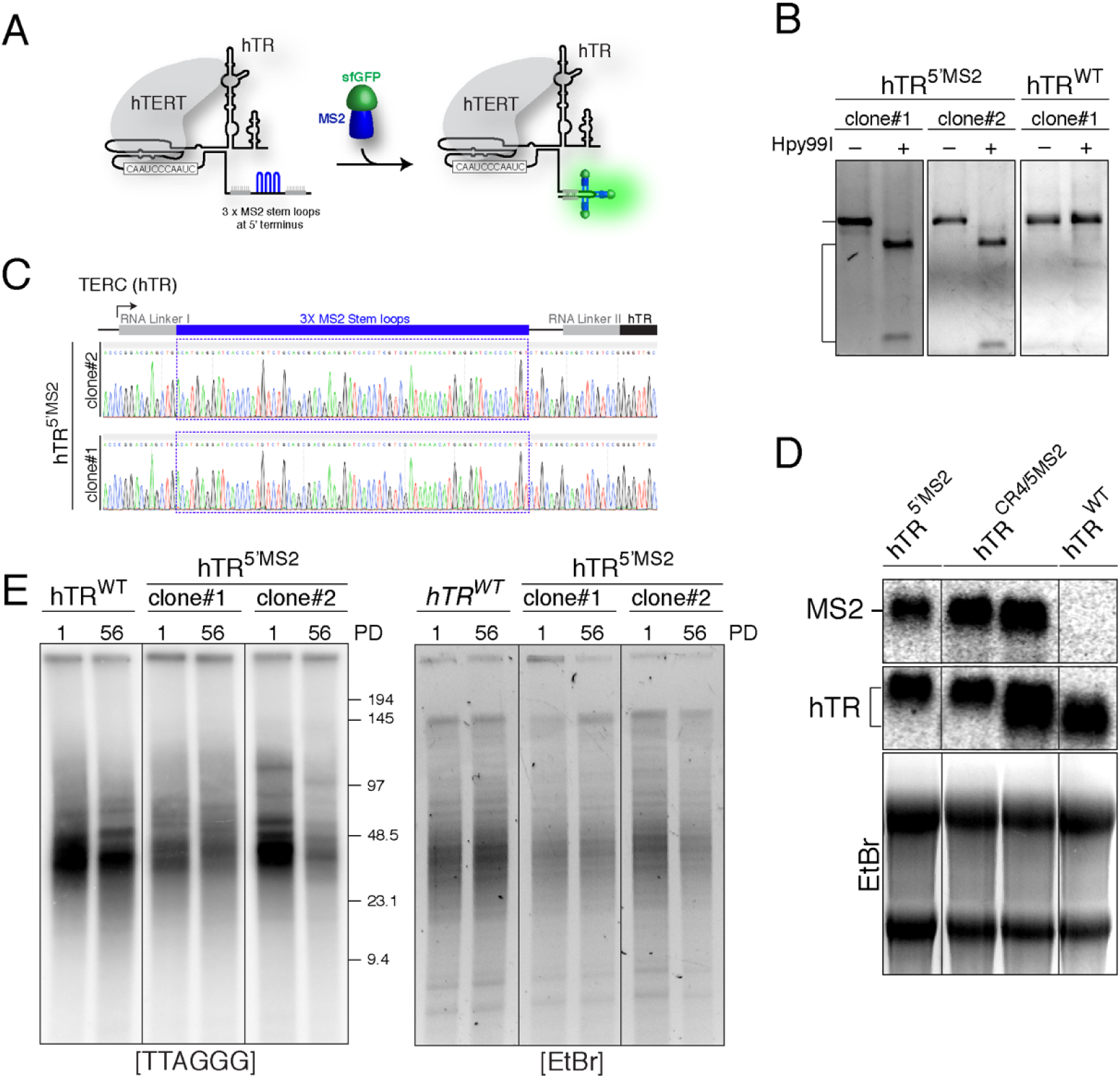
Tagging endogenous hTR with MS2 stem loops. **(A)** Schematic depiction of hTR tagged at the 5’-end with three MS2 stem-loops flanked by stabilizing RNA linkers. MS2-hTR is bound by GFP-labelled MS2 coat protein (GFP-MCP). **(B)** Genotyping PCR on DNA isolated from HeLa 1.3 cells to detect homozygous targeting of the endogenous hTR locus with three MS2 stem loops. Integration of the stem-loops generates a new Hpy99l cleavage site that is used to assess proper gene targeting. **(C)** Sanger sequencing to confirm proper integration of three stem-loops at the 5’-end of the hTR locus. **(D)** Northern blot analysis using MS2 (top) and hTR (bottom) probes on RNA isolated from the indicated cell lines. EtBr marking 28S and 18S rRNA serves as a loading control. **(E)** Southern blot analysis to monitor telomere length over time in HeLa 1.3 cells with the indicated genotype over time. *Mbo*I and *Alu*I digested DNA resolved on a gel electrophoresis and probed with end-labeled [AACCCT]_4_. EtBr-stained DNA serves a loading control.

We then performed CRISPR-Cas9 gene editing and modified the endogenous hTR locus in HeLa 1.3, a subclonal derivative of HeLa with long telomeres (Takai et al., 2010). We introduced three MS2 stem-loops at the 5’ end and the CR4/5 domain of hTR, independently. Genotyping PCR and Sanger sequencing confirmed proper gene targeting (Figures 1B-C and S1H) and Northern blot analysis indicated that MS2-tagged hTR was expressed at comparable levels as non-targeted cells (Figure 1D). Lastly, we assessed telomere length using Southern blot analysis and found that hTR modification did not alter telomere maintenance in *hTR^5’MS2^* and *hTR^Cr4/5MS2^* cells (Figures 1E and S1I).

### Single particle tracking identifies two populations of hTR with distinct dynamic properties

We transduced *hTR^5’MS2^* and *hTR^WT^* cells with lentiviral particles expressing GFP-MCP. Image acquisition with a spinning disk confocal microscope allowed the detection of single hTR particles in the nucleus of *hTR^5’MS2^* cells, but not in control *hTR^WT^* cells (Figure 2A). Particle tracking with the Trackmate algorithm (Tinevez et al., 2017) showed that the majority of telomerase RNA rapidly diffuse in the nucleoplasm and probe a large nuclear surface (Figure S2A and Movies S1-S3). Frequency distribution analysis identified a highly diffusive (mean logD: 0.09 µm^2^/s) fraction as well as a less mobile pool (mean logD: -0.69 µm^2^/s) (Figure 2B). We also detected MS2-hTR particles in the nucleolus and, interestingly, they displayed reduced diffusion rate compared to nucleoplasmic particles (Figure S2B-C and Movie S4). To better characterize the distinct populations of diffusive hTR molecules in *hTR^5’MS2^* cells, we overexpressed hTERT and hTERT-K78E, a mutant that disrupts telomerase interaction with TPP1 and prevents it recruitment to telomeres (Figure S2D)(Schmidt et al., 2014). We analyzed the distribution of diffusion coefficients obtained from MS2-hTR tracks and noted an enrichment in the fraction of slow diffusing particles in the presence of excess wild-type hTERT (Figure 2A-B). On the other hand, overexpression of hTERT-K78E resulted in a reduction in the percentage of less mobile hTR particles (Figure 2A-B). We therefore conclude that hTR particles with reduced mobility likely include telomere-bound telomerase. In contrast, rapidly diffusing particles potentially reflect telomere-unbound telomerase or free hTR molecules. We next sought to explore the localization and behaviors of these bound states.

**Figure 2.**
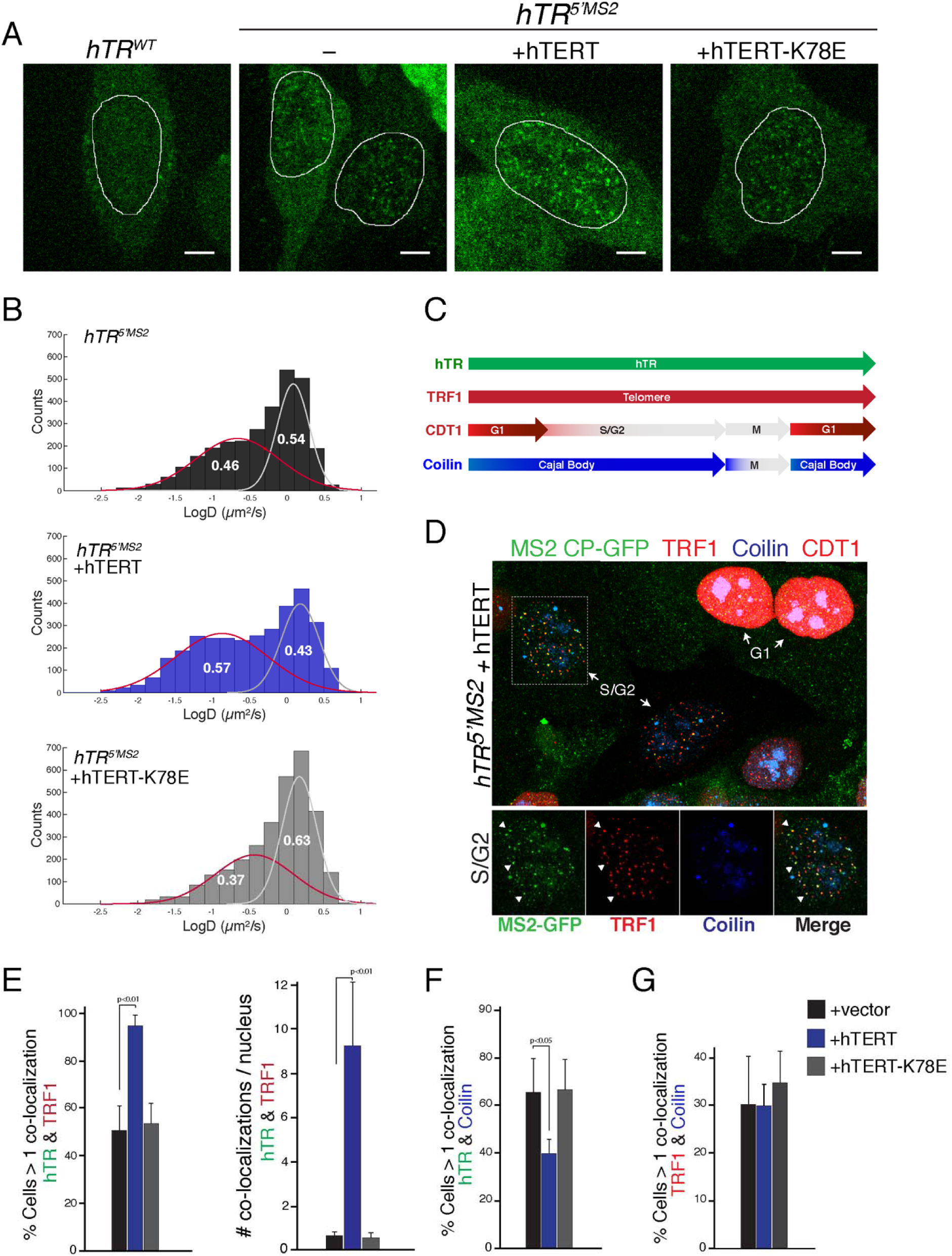
Single particle tracking of endogenous MS-tagged hTR displays two dynamic populations. **(A)** Still images from movies showing MS2-hTR foci (in green) in *hTR^5’MS2^* cells and *hTR^5’MS2^* cells overexpressing hTERT-WT or hTERT-K78E following transduction with GFP-MCP. Non-targeted (*hTR^WT^*) cells transduced with GFP-MCP serve as a negative control (scale bar = 5µm). **(B)** Frequency distribution analysis of diffusion coefficient from MS2-hTR particles in hTR^5’MS2^ cells overexpressing hTERT-WT, hTERT-K78E and control cells (3277 particles for hTR^5’MS2^+hTERT, 2875 particles for hTR^5’MS2^+hTERT-K78E and 2919 particles for hTR^5’MS2^). Gaussian fitting identified two mobile populations: a highly diffusive fraction (white curve) and a more static fraction (red curve). Percentage of each fraction is indicated. **(C)** Schematic representation of the different fluorescent markers to track MS2-hTR (GFP-MCP), telomeres (TRF1), and CBs (Coilin) and to mark the cell cycle (CDT1). **(D)** Colocalization analysis between MS2-hTR/GFP-MCP, BFP-Coilin and mCherry-TRF1 in hTR^5’MS2^ cells overexpressing hTERT. Marked G1 cells (top right) are mCherry-CDT1 positive while S/G2 cells are negative for mCherry-CDT1. Telomeres (labeled with mCherry-TRF1) are clearly visible in S/G2 cells and colocalization with MS2-hTR are indicted by arrowheads. Images were collected used a wide-field fluorescence laser scan confocal microscope (Nikon-Ti2 microscope). **(E)** Quantification of colocalization events between hTR and TRF1, hTR and coilin **(F)** and TRF1 and coilin **(G)**. Imaging was performed in *hTR^5’MS2^* cells overexpressing hTERT-WT, hTERT-K78E and control (vector only). Colocalizations are expressed as percentage of cells with more than 1 colocalization and as number of colocalization per nucleus. N=3 independent experiments with SDs and P values.

### Analysis of still images demonstrate that telomeres are not extended in CBs

We examined hTR localization at two important compartments for telomere homeostasis, the telomere itself and CBs where telomerase processing, assembly, and maturation take place. To visualize telomeres, we transduced *hTR^5’MS2^* cells expressing GFP-MCP with mCherry-tagged TRF1 and marked CBs with BFP-tagged coilin. In addition, we labelled *hTR^5’MS2^* cells with mCherry-CDT1 that is highly expressed in G1 cells but is rapidly degraded in S phase, when telomeres are extended. This was necessary to focus our image acquisition to this particular phase of the cell cycle during which telomerase extends telomeres (Figure 2C) (Tomlinson et al., 2006). Degradation of pan-nuclear CDT1 in S phase enabled the visualization of punctate mCherry at TRF1-bound telomeres (Figure 2D). Timelapse imaging with z-stack acquisition revealed that ∼40% of S phase cells display one or more colocalization event between hTR and TRF1. Furthermore, the association of telomerase with telomeres was enhanced following the overexpression of the catalytic subunit, but not in the presence of hTERT-K78E (Figure 2E). These results confirm that the TPP1-TERT interaction is essential for telomerase recruitment to telomeres.

Previous studies based on IF-FISH experiments reported the relocalization of replicating telomeres to the proximity of CBs (Jady et al., 2004; Jady et al., 2006). This led to a “hand over” model, whereby telomere extension took place within these nuclear bodies. We observed a reduction in MS2-hTR localization to CBs upon hTERT overexpression (Figure 2F), highly indicative of hTR eviction from CBs when telomeres are elongated. Furthermore, hTERT overexpression did not enhance the association of TRF1 with BFP-Coilin (Figure 2G). In summary, our data do not support the hand-over model and establish that productive interactions between telomerase and telomeres do not take place within CBs.

### Dynamic exchange of hTR between Cajal bodies and the nucleoplasm

In order to investigate the dynamic exchange of hTR molecules between the nucleoplasm and CBs, we co-expressed GFP-MCP and mCherry-labelled Coilin in *hTR^5’MS2^* cells and performed live-cell imaging. The majority of MS2-hTR freely diffused throughout the nucleoplasm and only a small fraction of hTR particles localized to CBs (Figure 3A and Movie S4). This observation is in disagreement with previous FISH experiments where hTR appeared to be predominantly concentrated in CBs (Tomlinson et al., 2006; Zhu et al., 2004). To corroborate our results, we performed dual IF-FISH analysis using Structured Illumination Microscopy (SIM). We adapted single molecule inexpensive FISH (smiFISH) to detect endogenous hTR with a set of 15 hTR-specific probes (Tsanov et al., 2016) and co-stained CBs with coilin antibody. SmiFISH-labelled hTR foci co-localized with CBs and at telomeres in hTR positive cells, including hTR^5’MS2^, hTR^Cr4/5MS2^ and the parental HeLa 1.3 cells (Figure S3A-B). SIM analysis validated live-cell imaging data and demonstrated that the majority of telomerase RNA molecules are distributed throughout the nucleoplasm with less than 10% of hTR accumulating in CBs (Figures 3B-C and S3C). Interestingly, SIM revealed that hTR foci accumulated at the periphery of CBs, as measured by the eccentricity ratio (Figure S3D).

**Figure 3.**
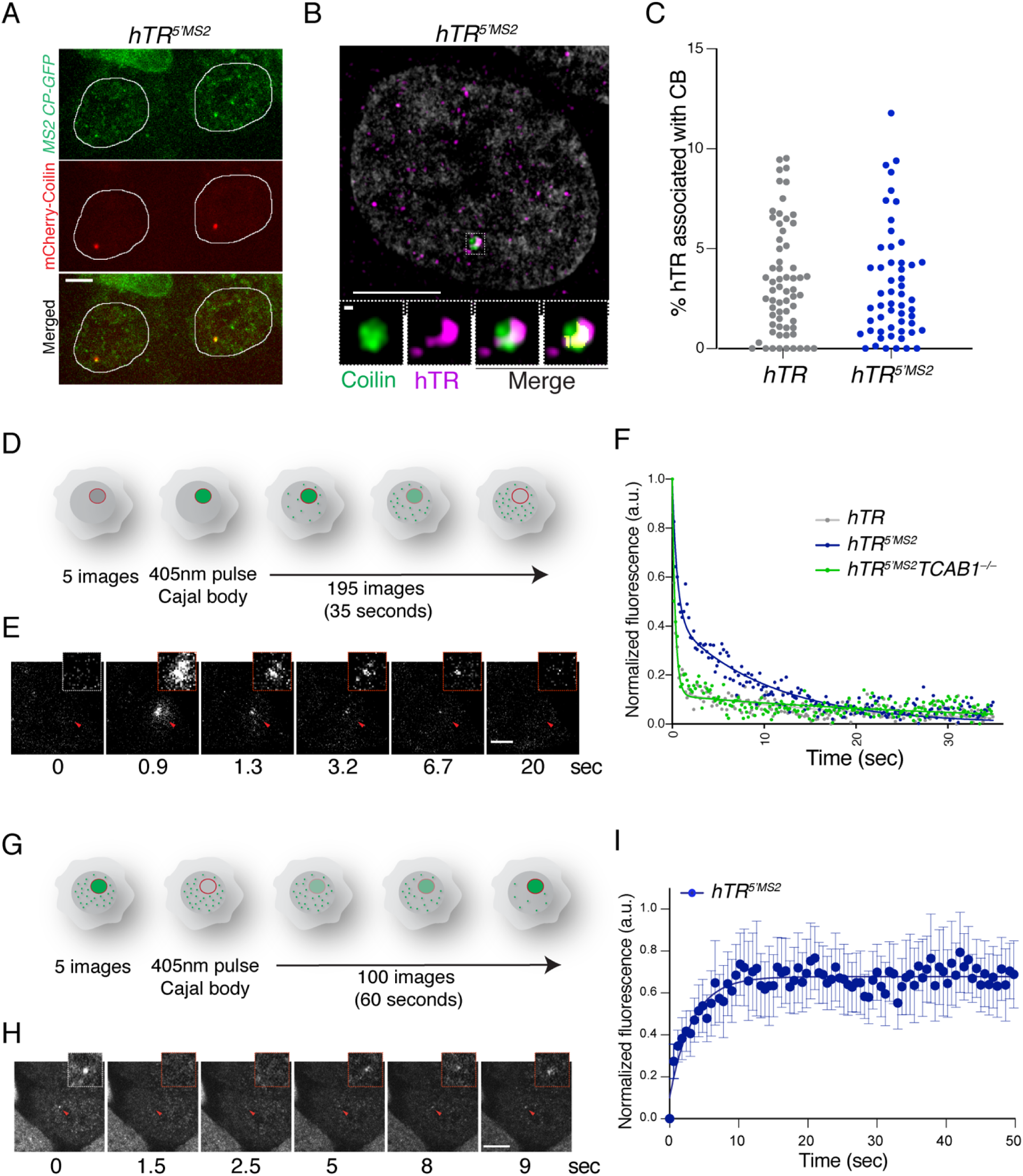
Dynamics of hTR association with Cajal bodies. **(A)** Still images obtained from movies which show colocalization between GFP-tagged MS2-hTR and mCherry-Coilin in *hTR^5’MS2^* cells. **(B)** Validation of MS2-hTR localization at CBs by combining smiFISH for hTR with Coilin immunofluorescence using super-resolution microscopy (SIM). Coilin-hTR colocalization regions are overlaid in yellow. **(C)** Quantification of hTR associated with CBs relative to the total amount of hTR per nucleus in *hTR^WT^* and *hTR^5’MS2^* cells. **(D)** Schematic depiction of photoactivation experiment. **(E)** Time-lapse of MCP-paGFP photoactivation in the Cajal body of a *hTR^5’MS2^* cell. Red arrow indicates the photoactivated Cajal body. **(F)** Fluorescence decay curves of MCP-paGFP following photoactivation in CBs from *hTR^WT^*, *hTR^5’MS2^* and *hTR^5’MS2^ TCAB1^-/-^* cells. For each cell line, four decay curves are averaged. **(G)** Schematic depiction of fluorescence recovery after photobleaching (FRAP) experiments. **(H)** Representative FRAP image from *hTR^5’MS2^* cells. Red arrow highlights photobleached Cajal body. **(I)** Graph depicting normalized fluorescence intensity recovery of photobleached MS2-hTR in CBs of *hTR^5’MS2^* cells as a function of time (N= 37 cells). All scale bars = 5µm, except Coilin scale bar = 2µm.

The MS2-hTR trajectories within CBs appeared less diffusive than nucleoplasmic hTR (mean logD of -0.97 µm^2^/sec *vs.* 0.09 µm^2^/sec; Figure 3A and Movie S4), prompting us to further characterize the properties of the RNA in these subnuclear structures. To that end, we fused a photoactivatable version of GFP with MS2CP (termed paGFP-MCP) and co-expressed the fusion protein along with mCherry-Coilin in *hTR^5’MS2^* cells. Specific photoactivation of paGFP-MCP in single CBs was followed by continuous acquisition of fluorescently labeled MS2-hTR (Figure 3D-E). In the absence of MS2-hTR, photoactivated paGFP-MCP displayed a rapid decay of fluorescence intensity, with a median half-life of 0.4 sec (Figure 3F). On the other hand, we observed two populations of MCP-paGFP in CBs of *hTR^5’MS2^* cells, including a rapidly diffusing fraction (median half-life of 0.6 sec) due to free MCP-paGFP, and a slow population (median half-life of 7.5 sec) indicative of MS2-hTR molecules (Figure 3F and Movie S5). To further validate that the paGFP-MCP signal was dependent on the presence of MS2-hTR in CBs, we repeated the photoactivation experiments in *hTR^5’MS2^ TCAB1^-/-^* cells generated using CRISPR/Cas9-based genome editing (Figure S3E-H). In the absence of TCAB1, fluorescence decay and MS2-hTR half-life in CBs were indistinguishable from that observed in control *hTR^WT^* cells (Figure 3F).

Having determined the half-life of hTR within CBs, we then used fluorescence recovery after photobleaching (FRAP) to explore the recruitment of MS2-hTR to CBs. Photobleaching of GFP-MCP was performed directly in CBs using a pulse of the 405 nm laser and followed by monitoring MS2-hTR recovery (Figure 3G-H and Movie S6). Fitting of a one-phase recovery curve to the average of these cells revealed that hTR recovers with a half-life of 2.3 seconds (Figure 3I). Taken together, our results indicate that a small subset of hTR molecules are targeted to CBs with a rapid on-rate and in a TCAB1-dependent manner. A slower off-rate leads to the accumulation of hTR in these nuclear bodies.

### hTERT facilitates hTR exit from Cajal bodies

To explore how hTR might be gated between CBs and telomeres, we explored the role of hTERT in this process. We generated hTERT knockout cells with CRISPR/Cas9 gene editing and derived two independent clones of *hTR^5’MS2^ hTERT^-/-^* cells that lack telomerase activity (Figure 4A-B). Using live-cell imaging and smiFISH, we noted a slight reduction in the accumulation of hTR in CBs in *hTERT^-/-^* cells (Figure S4A-B), suggesting that the loss of catalytic subunit does not block the recruitment of telomerase RNA to CBs. We then performed photoactivation experiments and observed a doubling of the half-life of MS2-hTR in CBs in two independent *hTR^5’MS2^ hTERT^-/-^* clones (Figure 4C-D). Complementation of *hTR^5’MS2^ hTERT^-/-^* cells with exogenous hTERT rescued hTR trafficking out of CBs (Figure 4D). Interestingly, FRAP experiments did not reveal a significant difference of hTR recovery dynamics in cells overexpressing hTERT relative to cells with endogenous hTERT (Figure S4C). Taken together, our data support a model in which the catalytic subunit of telomerase is important to drive the exit of hTR from CBs, but appears dispensable for targeting the RNA to these specialized subnuclear structures.

**Figure 4:**
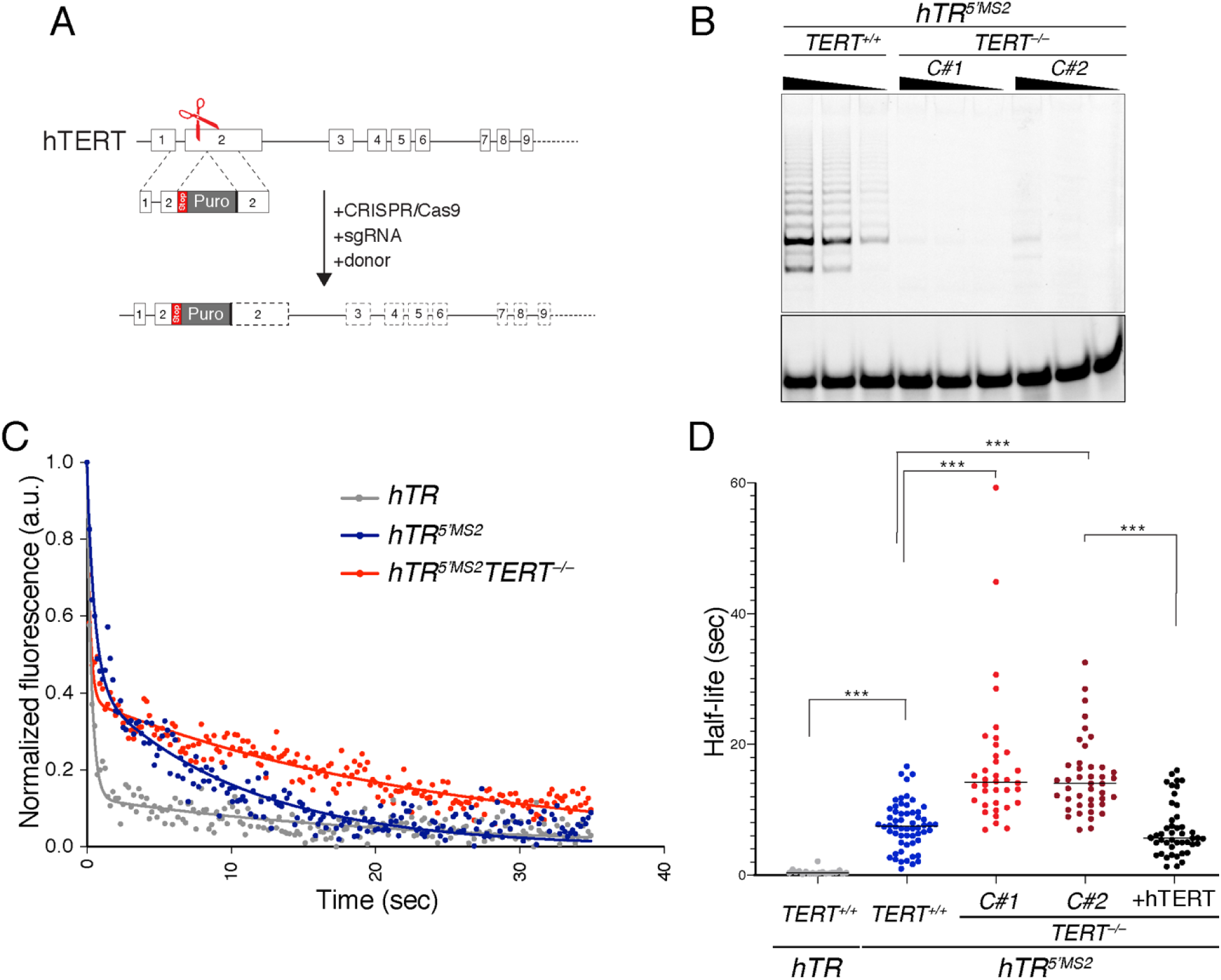
hTERT promotes the exit of hTR from Cajal bodies. **(A)** Genome editing strategy to knockout hTERT in *hTR^5’MS2^* cells. Cas9-mediated disruption of exon 2 of human *TERT* and the integration of a PURO selection cassette. **(B)** Telomere Repeat Amplification Protocol (TRAP) assay to validate the loss of telomerase activity in two clonally derived *hTR^5’MS2^ TERT^-/-^* cells lines. **(C)** Fluorescence decay of MCP-paGFP following photoactivation in CBs from *hTR^WT^*, *hTR^5’MS2^*, and *hTR^5’MS2^ TERT^-/-^* cells. **(D)** Half-life of MS2-hTR in CBs of cells with the indicated genotype and treatment (N=2 independent experiments). *** p< 0.005.

### Multiple dynamic interactions govern telomerase-telomere association

We turned our attention to the association of telomerase RNA with its substrate – the telomere – and performed live-cell imaging of MS2-hTR and mCherry-TRF1 in S phase *hTR^5’MS2^* cells (Figure 5A). Single-particle tracking was achieved by constant acquisition at 100 msec/frame for a total of 10 sec and revealed that within this timeframe, hTR particles colocalized with up to 50% of visible telomeres per cell (Figure 5B and Movie S7). As expected, overexpression of hTERT increased the fraction of hTR-bound telomeres while hTERT-K78E diminished the co-localization of MS2-hTR and mCherry-TRF1 (median of 31% and <1%, respectively; Figure 5B). Analysis of hTR trajectories at telomeres highlighted a set of localized tracks that colocalized with TRF1 for more than 5 frames and were enriched in the presence of hTERT but not hTERT-K78E (Figure S5A-B). Within the 10 second image-acquisitions, the median residence time of hTR at telomere in hTR^5’MS2^ cells was ∼0.7 seconds (Figure 5C). Disrupting the interaction between TERT and TPP1 greatly diminished the frequency of telomere-hTR colocalization, with a few events remaining within the same dwell time. In contrast, overexpression of hTERT led to a notable increase in colocalization events with longer residence time, including several tracks that lasted well-beyond the 10 sec movies (Figure 5C). To better study the dynamic of these long interactions, we used time intervals of 5 seconds for a total of 300 seconds acquisition. While long lasting interactions were observed in hTR^5’MS2^ cells, their frequency was more pronounced in cells overexpressing hTERT and ranged between 10 and 220 seconds with a median of 25 sec (Figure 5D). Fitting the distribution of hTR dwell time at telomeres in *hTR^5’MS2^* and *hTR^5’MS2^* +hTERT cell lines revealed similar off-rate, half-life, and mean residence time (Figure S5C-D), suggesting that increased hTERT levels did not alter the dynamics properties of telomerase at telomeres. Notably, visualization of hTR tagged with 3MS2 stem-loops at the CR4/5 domain yielded a similar dynamic profile to that of 5’MS2-hTR (Figure S5J).

**Figure 5:**
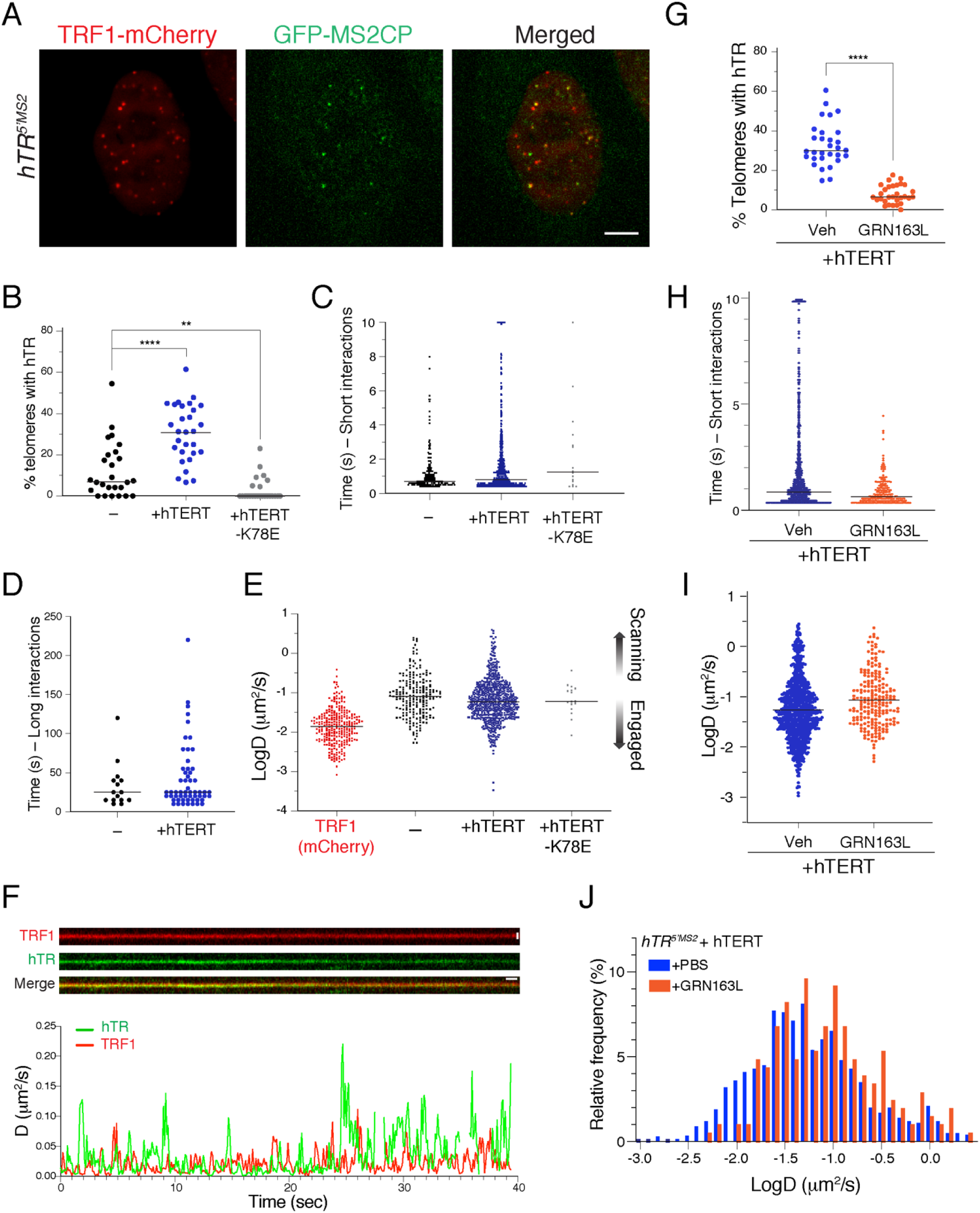
Deciphering the dynamic properties of hTR interaction with telomeres. **(A)** Representative image depicting the colocalization of MS2-hTR/MCP-GFP with telomeres labelled with mCherry-TRF1 in *hTR^5’MS2^* (Scale bar = 5µm). **(B)** Percentage of telomeres colocalizing with MS2-hTR in *hTR^5’MS2^* cells and in cells overexpressing hTERT-WT or hTERT-K78E. Median values are 7% for hTR^5’MS2^, 31% for hTR^5’MS2^ + hTERT WT and <1% for hTR^5’MS2^ + hTERT-K78E. ** p< 0.01; **** p< 0.001. **(C)** Short-term residence time of hTR particles at telomeres in HeLa *hTR^5’MS2^* cells, and in *hTR^5’MS2^* cells overexpressing hTERT-WT and hTERT-K78E. **(D)** Residence time of long interactions between hTR particles and telomeres in *hTR^5’MS2^* with or without hTERT overexpression. **(E)** Graph depicting diffusion coefficients of hTR particles at telomeres in *hTR^5’MS2^* cells, and in *hTR^5’MS2^* cells overexpressing hTERT-WT and hTERT-K78E. LogD of TRF1 was used as a reference for telomere mobility. Top and down arrows correspond to scanning and engaged hTR particles, respectively. **(F)** Dual-camera imaging of hTR at a telomere. Top – representative kymograph for hTR and TRF1 tracks. (Vertical scale bar = 0.80 µm, horizontal scale bar = 1 second). Bottom – running-window analysis of diffusion coefficients for TRF1 and hTR as a function of time over a span of 40 seconds. **(G)** Percentage of telomeres associated with MS2-hTR in *hTR^5’MS2^* cells treated with 3µM GRN163L for 24 hours. **** p< 0.001. **(H)** Graph depicting residence time of MS2-hTR at telomeres in cells treated with vehicle or GRN163L. **(I)** LogD analysis of *hTR^5’MS2^* +hTERT cells treated with GRN163L. Note the reduction in the frequency of slow diffusive particles upon treatment with 3 µM GRN163L for 24 hours. **(J)** Histogram depicting the relative frequency of LogD distribution for MS2-hTR in cells as in (**I**).

### Telomerase oscillates between scanning and engaged states at telomeres

Analysis of hTR trajectories throughout the nucleoplasm highlighted a small fraction of RNA particles with limited diffusion (Figure 2A-B), which we postulated to be constrained by telomere-interactions. Therefore, we set out to investigate the diffusive properties of MS2-hTR particles that co-localize with their substrate. We first assessed telomere mobility and determined the diffusion coefficient (D) of TRF1, which displayed a median logD of -1.855 µm^2^/s with the fastest molecule exhibiting a D∼0.40 µm^2^/s (Dimitrova et al., 2008; Schmidt et al., 2016). We determined the diffusion coefficient of hTR molecules confined at telomeres and compared it with TRF1 diffusion. Interestingly, a significant fraction of hTR particles were more diffusive than the telomere itself, even when confined to their substrate (Figure 5E). This suggest that following TPP1-dependent recruitment to telomeres, telomerase display two diffusive states, which we term “scanning” and “engaged”. hTR particles with a faster diffusion rate than the telomere chromatin reflects scanning interactions. On the other hand, particles with similar diffusion properties as telomeres are engaged with their substrate. To validate these results, we performed dual-camera image acquisition and tracked MS2-hTR and TRF1-bound telomeres simultaneously. Kymographs showed that colocalization between an hTR particle and a telomere could be traced for several seconds (Figures 5F and S5E, and Movie S8). Using a running-window analysis of the diffusion coefficient, we noted periods in which hTR and TRF1 displayed similar diffusion rates, but also instances where hTR had faster diffusion coefficient than TRF1. Similar results were obtained upon measuring the radius of gyration of hTR and telomere (Figure S5F-G). In summary, a given hTR molecule is capable of alternating between scanning and engaged modes when interacting with the telomere.

### Telomerase retention is dependent on RNA-DNA base pairing

So far, our data indicate that hTR interaction with telomeres oscillates between scanning and engaged modes, with a dwell-time ranging between 0.5 and 220 seconds. We hypothesized that engaged interactions reflect a retention step during telomere elongation where base pairing of hTR template with the single-stranded overhang restricts telomerase mobility and increases its residence time at chromosome ends. To better define telomerase retention, we treated hTR^5’MS2^ cells overexpressing hTERT with GRN163L (Imetelstat) that blocks the template region of telomerase and thus inhibits hTR-ssDNA interaction at telomeres (Herbert et al., 2005). In the presence of GRN163L, colocalization between MS2-hTR and telomeres was significantly reduced (Figure 5G) and its dwell time did not exceed 5 seconds (Figure 5H). Survival probability analysis revealed a four-fold reduction in the half-life of hTR at telomeres and a concomitant increase in its off-rate when base-pairing with the telomere overhang was blocked (Figure S5H-I). Interestingly, the population of telomere-localized hTR with the lowest diffusion coefficient was nearly eliminated in the presence of GRN163L (Figure 5I-J). Altogether, our results suggest that following TPP1-mediated recruitment, telomerase retention at telomeres is an independent step driven by base-pairing interactions between the template region of hTR and telomere ssDNA.

### ATM acts upstream of TPP1 to facilitate telomeres access to telomerase

We next aimed to decipher the different states of telomerase-telomere interactions using genetic perturbations known to alter telomere length homeostasis. Signaling by ATM and ATR was previously shown to promote telomere elongation, however, the exact step(s) of telomerase trafficking controlled by these kinases remain unknown (Lee et al., 2015; Tong et al., 2015). We treated *hTR^5’MS2^* cells overexpressing hTERT with the kinase inhibitors KU60019 and VE-822 at concentrations previously shown to inhibit the activity of ATM and ATR, respectively (Fokas et al., 2014; Hickson et al., 2004). We then monitored the dynamic properties of hTR trajectories at telomeres using live-cell imaging. Consistent with previous studies, ATM inhibition led to a decrease in the percentage of telomerase at chromosome ends (Figure 6A). In contrast, inhibition of ATR kinase did not significantly impair telomere-telomerase association (Figure 6B). The inhibition of ATM activity did not impact the duration of short interactions (Figure S6A), nor did it significantly alter long-lasting interactions (Figure S6B). We confirmed these results using shRNA-mediated depletion of ATM (Figure 6C and S6D-F). Collectively, our data demonstrate that while ATR is dispensable for telomerase association with chromosome ends, ATM acts upstream and increases the frequency of telomerase recruitment without impacting its dwell time at telomeres.

**Figure 6:**
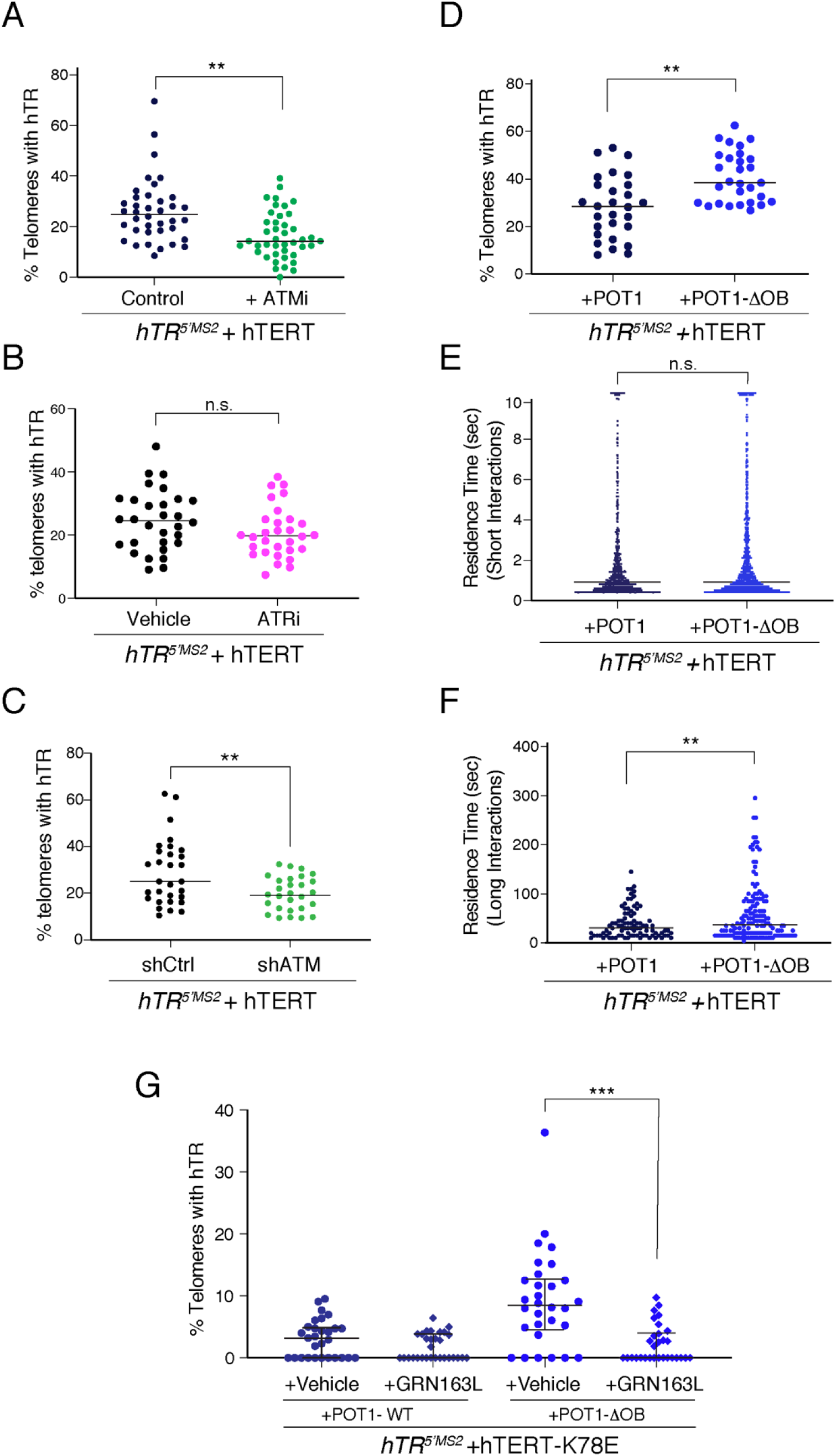
ATM and POT1 regulate different steps of hTR interaction with telomeres. **(A)** Treatment of hTR^5’MS2^+hTERT cells with ATM inhibitor reduces the colocalization between hTR and telomeres. Cells were treated with 3 µM of KU60019 (ATMi) and DMSO (control) for 24 hrs. ** p<0.01. (N=42 cells with ATMi and 38 DMSO-treated cells). **(B)** Percentage of telomeres colocalized with hTR in hTR^5’MS2^ +hTERT cells treated with DMSO (control) and 0.5 µM of VE822 (ATRi). (N=30 cells). **(C)** shRNA-mediated depletion of ATM reduced the localization of hTR at telomeres. Percentage of telomeres colocalization with hTR in hTR^5’MS2^ +hTERT cells expressing shCtrl or ShATM. ** p<0.01. (N=30 cells). **(D)** Expression of mutant POT1 increases the localization of hTR at telomeres. Graph represents percentage co-localization of TRF1 and MS2-hTR in hTR^5’MS2^+hTERT cells transduced with POT1-WT and POT1-ΔOB. (N=29 cells expressing POT1-ΔOB and 28 cells expressing POT1-WT). ** p<0.01. **(E)** Short-term residence time of hTR at telomeres is independent of POT1 (N= 29 cells for POT1-ΔOB and 28 for POT1-WT). **(F)** Long-term residence time of hTR at telomeres is increased in hTR^5’MS2^ cells expressing POT1-ΔOB compared to POT1-WT. **p<0.01. (N=10 cells). **(G)** POT1 inhibits the interaction of telomerase RNA template with single-stranded telomere overhang. hTR^5’MS2^ +hTERT-K78E cells expressing POT1 WT or POT1-ΔOB were treated 24h with GRN163L or vehicle control. (N=30 cells per condition). ***p<0.005.

### POT1 negatively regulates telomere length by preventing telomerase retention at chromosome ends

It is well established that the shelterin component, POT1 acts as a negative regulator of telomerase. Expression of POT1 mutations, including those commonly found in human cancers and those lacking the OB fold domain of POT1 (POT1-ΔOB), lead to rapid telomere elongation in cancer cells (Loayza and De Lange, 2003; Pinzaru et al., 2016). However, FISH- and ChIP-based assays failed to detect the impact of POT1 depletion or mutations on the telomerase interaction with telomeres *in vivo* (Abreu et al., 2010; Gu et al., 2017). We transduced *hTR^5’MS2^* cells overexpressing hTERT with lentiviral particles expressing POT1-WT and POT1-ΔOB (Figure S6G-H) and observed an increase in the percentage of hTR colocalization with telomeres in the presence of mutant POT1 (Figure 6D). Notably, POT1-ΔOB had no effect on short-term interactions (Figure 6E), instead, it enhanced long-interactions of telomerase with its substrate (Figure 6F). In an independent set of experiments, we transduced *hTR^5’MS2^* cells expressing POT1-WT and POT1-ΔOB with lentiviral particles for hTERT-K78E. As expected, the localization of hTR to telomeres was greatly diminished in cells co-expressing hTERT-K78E and POT1-WT (Figures 6G and S6I). However, we detected significant association of hTR with telomeres in the presence of mutant POT1, even when the interaction between hTERT and TPP1 was reduced (Figure 6G and S6I). This observation suggested that enhanced telomere-telomerase interactions observed in the presence of POT1-ΔOB are mediated by hTR base-pairing with telomeric ssDNA. Consistent with this idea, hTR-telomere interactions were diminished upon treatment of *hTR^5’MS2^* cells co-expressing POT1-ΔOB and hTERT-K78E with GRN163L (Figure 6G). Altogether, our data support a model where POT1 acts downstream of TPP1 and inhibits telomerase retention by occluding the single-stranded telomere overhang from binding to the template region of hTR.

## DISCUSSION

### A “Recruitment – Retention” model for telomere extension

Since its discovery over 30 years ago (Greider and Blackburn, 1985), telomerase has been extensively studied through biochemical and genetic analysis in different model organisms. However, the dynamic properties of this enzyme *in vivo* are poorly understood. Here, we successfully adapted the MS2 tagging system and fully depicted the spatio-temporal regulation of the specialized RNA throughout the various steps of telomerase assembly, maturation, trafficking, and recruitment to telomere ends. Our results reveal a two-step, “Recruitment – Retention” model (Figure 7), where TPP1-mediated recruitment leads to a series of short and highly diffusive telomere-telomerase associations. Subsequently, a subset of telomerase molecules is retained at telomeres through base-pairing of the hTR template region with the single-stranded telomere overhang. The latter molecules exhibit a longer residence time and slow diffusive properties. Whether telomerase retention is enacted by factors along the telomere chromatin is yet to be determined. It is worth noting that the retention step has not been delineated for telomerase in lower eukaryotes. Emerging evidence from yeast indicate that telomerase recruitment is driven by protein-protein interactions in a way that is analogous to TPP1-mediated recruitment of hTERT (Chandra et al., 2001; Tomita and Cooper, 2008). Interestingly, genetic studies identified specific alleles, including Est3 in budding yeast (Rao et al., 2014) and TAZ1 K57A in fission yeast (Armstrong et al., 2014) that support telomerase recruitment but fail to maintain telomere length. It has been postulated that these mutants are defective in activating telomerase. Nevertheless, it is tempting to speculate that such alleles modulate a step equivalent to telomerase retention.

**Figure 7:**
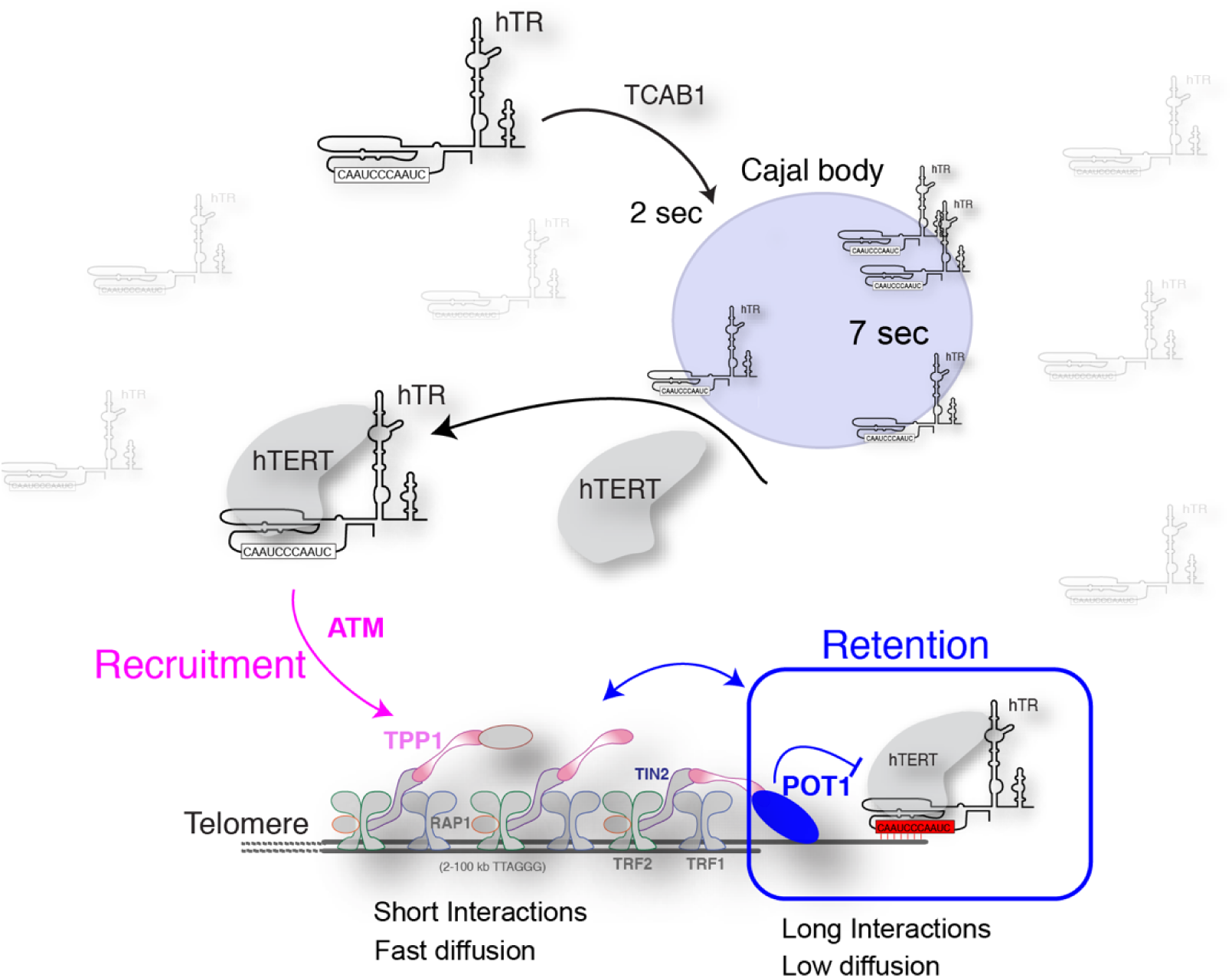
Model for hTR dynamics between CBs, nucleoplasm, and telomeres. Our study unveils the life-cycle of hTR in cancer cells. The majority of hTR particles are highly diffusive throughout the nucleoplasm. A small fraction of the RNA is targeted by TCAB1 to CBs, where it is retained for an extended time. The exit of hTR from CBs is facilitated by hTERT and the assembly of a full RNP. Telomere elongation is governed by a “Recruitment – Retention” model. Recruitment of telomerase to telomere is mediated by interaction between hTERT and TPP1, and facilitated by ATM activity. This allows a series of short interactions in which telomerase is more diffusive than its substrate. A fraction of these telomerase particles is retained at chromosome ends upon base pairing of template region of hTR with the telomere overhang, Interestingly, the retention step is regulated by the binding of POT1 OB-fold domains to the telomeric ssDNA.

### Cajal bodies as assembly sites for telomerase RNP

hTR was first reported to accumulate to CBs in cancer cells and the organelle was later shown to be critical for hTR biogenesis and maturation (Schmidt and Cech, 2015). So far, investigation of telomerase association with CBs has been limited to the analysis of fixed cells using RNA FISH. This led to the current dogma that telomerase RNA is predominantly concentrated in CBs and telomere-targeting to these nuclear bodies facilitate their elongation (Tomlinson et al., 2006; Zhu et al., 2004). In contrast, data from live-cell and super-resolution imaging show that less than 10% of hTR molecules reside in CBs (Figure 3a-c). Furthermore, we demonstrate that telomere elongation takes place outside CBs (Figure 2f-g). hTR molecules have an extended residence time in these nuclear bodies (Figure 3d-f) and this could explain, at least in part, why low sensitive RNA FISH only detects endogenous hTR in CBs. Our data is consistent with a model where CBs act as temporary assembly sites for properly modified and processed hTR molecules to be accessed by hTERT. Unlike hTR, the association of hTERT with CBs is highly transient (Schmidt et al., 2016), implying that the loading of hTERT onto a hTR scaffold is among the last steps during telomerase assembly. Interestingly, super-resolution imaging revealed that hTR molecules protrude from the coilin core of CBs (see also (Jady et al., 2004)), suggesting a model where hTERT may assemble with hTR at the edge of CBs. What accounts for extended residence time of hTR in CBs is unknown. Coilin contains an intrinsically disordered RG-rich motif that has the propensity to oligomerize and create higher order structures that could potentially trap RNA within these nuclear bodies (Hebert et al., 2001). Alternatively, homotypic and heterotypic RNA-RNA interaction could confine hTR mobility in CBs.

### Unique properties of hTR relative to hTERT and yeast *TLC1*

The dynamic behavior of the reverse transcriptase was monitored using Halo-tag and demonstrated that hTERT is highly diffusive in the nucleoplasm and displays long and short interactions with telomeres (Schmidt et al., 2016). Overall, these observations are consistent with results obtained in our study, however, there are important differences between the dynamics of HaLo-hTERT and MS2-hTR. First, HaLo-hTERT interaction with Cajal bodies is more transient than hTR, potentially highlighting distinct maturation pathways for each subunit. Second, MS2-hTR particles are present in the nucleolus, whereas HaLo- and GFP-tagged hTERT are excluded from this subnuclear domain in cancer cells (Schmidt et al., 2016; Wong et al., 2002). Third, MS2-hTR and HaLo-hTERT behave differently at telomeres. For instance, using dual-camera imaging we observed scanning interactions between hTR and telomeres (Figure S5A-B). In contrast HaLo-hTERT interactions are either probing (transient) or stably associated (localized) at telomeres. Furthermore, while the kinetic of hTERT binding to telomeres was not affected by GRN163L treatments (Schmidt et al., 2018), the drug significantly impacted the dwell time and off-rate of MS2-hTR (Figure 5). The different dynamic properties could be explained by a faster rate of acquisition during imaging of HaLo-hTERT compared to MS2-hTR. Although, we cannot rule out that compromised telomerase activity due to hTERT tagging (Chiba et al., 2017; Schmidt et al., 2018) could have altered the dynamic behavior of the enzyme at telomeres.

In addition, our analysis of human telomerase RNA reveals significant differences in the dynamics of telomere elongation between human and budding yeast. We previously employed the MS2 system to label telomerase RNA in *S. cerevisiae* and found that telomere elongation in late S phase is dependent on stable accumulation of multiple telomerase complexes with no apparent scanning interactions (Gallardo et al., 2011). Unlike yeast *TLC1* RNA, hTR molecules did not cluster at human telomeres. Mutants affecting telomere elongation in yeast – most notably being tel1 kinase – inhibit telomerase recruitment at short telomeres and disrupt clustering of yeast telomerase (Gallardo et al., 2011; Goudsouzian et al., 2006). In the case of human telomerase, we found that inhibition of ATM, but not ATR, impacts the access of the RNP to telomeres. In both cases, downstream targets of ATM remain elusive. Furthermore, how these kinases influence telomerase targeting to short *vs.* long telomeres is unknown.

### POT1 regulates telomerase retention at telomeres

Our work also provides insight into the mechanism by which POT1 regulates telomerase activity at telomeres. So far, conflicting models have been proposed for the function of POT1 at telomeres. Genetic and biochemical evidence suggest that POT1 inhibits telomerase, possibly by sequestering the telomeric overhang. Specifically, deletion or point mutations in the OB-fold domain of POT1 lead to unrestricted telomere elongation by telomerase (Loayza and De Lange, 2003). However, depletion or mutations in POT1 did not reduce telomerase access to telomeres, as measured by RNA-FISH or ChIP (Abreu et al., 2010; Gu et al., 2017). Moreover, biochemical assays have shown that the TPP1-POT1 complex stimulates telomerase processivity *in vitro* (Wang et al., 2007). Our results uncover the mechanism by which POT1 regulate telomere elongation *in vivo*. We show that OB-fold mutation of POT1 enhances telomerase retention by increasing overhang accessibility. Our data support a model whereby POT1 OB-fold domains, where most cancer-associated mutations incur, compete with hTR for the binding to the telomere overhang. The negative regulation of telomerase retention by POT1 underscores the importance of restricting telomere elongation by telomerase and highlights a mechanism that has gone awry in an increasing number of solid and lymphoid malignancies with POT1 mutations.

## Acknowledgements

We thank Steven Artandi, Titia de Lange, and Jerry Shay for providing key reagents for this study. We acknowledge M. Anh-Tien Ton (Serohijos lab) and Dr. Nicolas Stifani (UdeM) for their assistance in writing scripts for image analysis. Frank Yeung, Michal Cammer, and Aleks Penev are thanked for their help in the early stages of the study. We acknowledge the microscopy core at NYU School of Medicine and UdeM. We thank members of the Sfeir and Chartrand lab for critical reading of the manuscript. This work was supported by grants from the NIH-NCI to A.S., the Canadian Institutes of Health Research (PJT-162156) and the Canadian Cancer Society Research Institute (CCSRI) to P.C. P.C holds a Research chair from the Fonds de Recherche du Québec-Santé (FRQS).

## Authors Contribution

A.S. and P.C. conceived the experimental design. H.L. performed key experiments with help from E.Q., D.G., E.P., and D.C. D.G. constructed and validated the MS2-hTR system. E.Q. and M.J.S. performed experiments related to CBs. D.C. performed still image analysis. H.C. performed all gene editing experiments. A.S. and P.C. wrote the manuscript and act as co-corresponding authors. All authors discussed the results and commented on the manuscript.

## Authors information

Correspondence and requests for materials should be addressed to P.C. and A.S.

## MATERIAL and METHODS

### Cell culture procedures and treatments

293T cells were obtained from (ATCC). HeLa 1.3 cells were cloned from parental HeLa and have long telomeres, identified by the de Lange lab (Takai et al., 2010). Cells were cultured in Dulbecco’s Modified Eagle Medium (DMEM, Corning) supplemented with 10% Fetal Bovine Serum (BCS, Gibco), 2 mM L-glutamine (Gibco), and 100 U/ml Penicillin-Streptomycin (Gibco). Cells were passaged every 48-72 hours and maintained mycoplasma free by using Plasmocin (Invivogen) per manufacturer indication. For ATM inhibition, cells were treated with 3 µM of KU60019 (Tocris Bioscience) for 24 hours. ATR inhibition was achieved by treating cells with 0.5 µM of VE822 (Selleck Chemicals) for 3 hours. For inhibition of Telomerase template annealing, 3 µM of GRN163L was added for 24 hours. Transduction of MS2-GFP, hTERT, and TCAB1, was achieved using lentivirus-containing supernatant generated in 293T cells. Viral supernatants were supplemented with 4 g/ml polybrene. Infections were followed by puromycin selection for 3 days or sorted with GFP in the case of MCP-GFP.

### CRISPR/Cas9 gene editing

#### hTR-MS2 targeting

HeLa MS2 kock-in cells were generated by CRISPR/Cas9 targeting. Briefly, HeLa cells were transfected using Lipofectamine 3000 (ThermoFisher) using a donor plasmid of 3xMS2 repeats flanked by homologous hTR sequences and 3 different gRNAs cloned into a Cas9 Nuclease plasmid, pSpCas9(BB) (PX330) (Addgene). Targeting was done for the 5’ and CR4/5 sites, independently. (gRNA sequences: gRNA4 (5’) TCCGCAACCCGGTGCGCTGC, gRNA5 (5’) GCAGCGCACCGGGTTGCGGA and gRNA9 (CR4/5) TCGGCCCGGGGCTTCTCCGG]. Cells were selected 24 hours post transfection using 600 ng/mL of Puromycin. Selection was removed after 24 hours and surviving cells were plated at clonal density. Individual clones were isolated and used to establish targeted cell lines.

The initial genotyping PCR was performed to determine donor integration using primers that bind within donor sequence. (RA_in Fw3: AGCCCGCCCGAGAGAGTG and LA_in Rv1: ATGTGTGAGCCGAGTCCTGGG). Hits identified from the initial genotyping screen were further validated by additional PCR and by Northern blot analysis. To validate site-specific integration of the MS2 sequences, a restriction enzyme digestion was performed on a PCR product obtained using hTR and donor primers (LA_out Rv2: AGAATGGCCTGTTTGTTTCTTTCAACCTAGTGGG and RA_in Fw2 TTTTTAAGGTAGTCGAGGTGAACCGCG). 1450bp amplicon corresponds to non-targeted allele, while a band of 1550bp indicates MS2 integration. Amplicon from targeted alleles yield two bands when digested with Hpy991. Successful targeting of the 5’ end of hTR generated bands of 1200bp and 350bp, while targeting at the Cr4/5 region 900bp and 600bps indicated successful targeting of the CR4/5 region. Finally, to confirm site the site-specific integration, the hTR regions were sequenced.

#### TCAB1 targeting

TCAB1 (WRAP53) was targeted in MS2_hTR HeLa cells (clone 15.5) using CRISPR/Cas9 targeting. A mixture of 2 sgRNAs that specific for exon 1 (∼100bp downstream of the ATG) were cloned into Cas9 nuclease (pX330_puro). gRNA sequence: GTCCGCATTTTTATTCATCG and TTTATTCATCGGGGAAGCGT. A single stranded DNA oligo to serve as a homologous recombination donor was designed and purchased from IDT. The donor introduces a premature stop codon, a frameshift substitution and XbaI restriction enzyme. HeLa cells were transfected with the gRNAs and ss donor DNA using a nucleofector (Lonza, solution SE, Program CN-114, 1M cells/reaction +8ug DNA). Puromycin (600ng/mL) was added 24 hours after transfection and cells were selected for 3 days. clonally derived cell lines were established. Clones were screened for ssDonor integration and TCAB1 KO by PCR amplification and XbaI restriction enzyme digest (primers: F4: AGAAGAGGAGGGAAGCACAGG and R2: TTCCATGGCTTTT CCAGACCCC). A wildtype band corresponds to 497bp and cleaved into 220bp and 227bp upon targeting. Proper integration was further confirmation by sequencing the locus.

#### hTERT targeting

hTERT was knocked out using CRISPR/Cas9 targeting of MS2_hTR HeLa1.3 cells (clone 15.5). gRNAs were designed to target exon 2 of the TERT locus. Two gRNAs were cloned into the EmptypSpCas9n(Bbs)(Bsa)-2A-GFP plasmid to allow for double nuclease cutting of the target sequence sites. [gRNA sequences: gRNA1: CACCGCTTCGGGGTCCACTAGCGTG and gRNA5:CACCGCGCAGCTACCTGCCCAACA]. A donor plasmid was designed and contained two 500bp sequenced with homology to the hTERT locus. Cells were transfected with the donor plasmid and gRNAs/Cas9 and sorted for GFP expression of the Cas9. Cells were recovered for 1 week and then selected for Puro integration. Surviving cells were plated for clonal expansion and screened for on-target Puro integration and hTERT knockout. (primers: geno Rev RAD out: AACAGGGGCCGCATTTGCCAGTAGC and genoPURO fwd: GGGTGCCCGCCTTCCTGG)

### Western Blot analysis

Cell pellets were collected and protein was isolated by lysing the cells using a RIPA buffer and sonication. Protein was quantified using a BCA Protein Assay Kit (ThermoFisher Pierce) and 20µg of protein/sample was loaded in the gel. the gel was transferred to a membrane using the BioRad turbo blot. The membrane was blocked in 5% milk for 1 hour and then incubated with an affinity purified TCAB1 antibody (Artandi Lab) diluted 1:10000 in 5% milk at 4°C overnight, or gamma-Tubulin, diluted 1: 5000 in 5% milk at 4°C, overnight. Primary antibody was washed 3 times with TBS-T and blotted with either rabbit (1:5000 - TCAB1) or mouse (1:5000 - Tubulin) for 30 minutes at room temperature. The membrane was washed again and the signal was detected using BioRad ECL. Recombinant anti-ATM antibody (Y170, Abcam ab32420), anti-c-Myc antibody clone 9E10 (Millipore, 1:1000 dilution), and Recombinant Anti-Telomerase reverse transcriptase antibody (Y182, Abcam ab32020) were incubated diluted 1:1000 in 3% BSA in TBS-T at 4°C, overnight).

### Telomere length analysis

Analysis of telomere length was performed using pulse-field gel electrophoresis and in gel hybridization as previously described (Hockemeyer et al., 2006).

### Northern Blot analysis

Total cellular RNA was prepared using RNeasy Mini Kit (Qiagen), according to the manufacturer instructions. 20 μg RNA was loaded onto 1.5% formaldehyde agarose gels and separated by gel electrophoresis for 17hours at 40V / 4°C. RNA was transferred to a Hybond membrane. The blot was prehybridized at 60°C for 1 h in Church mix (0.5 M Na_2_HPO_4_ (pH 7.2), 1 mM EDTA, 7% SDS, and 1% BSA), followed by hybridization at 60°C overnight with probes for hTR and MS2. A combination of six P^32^-labeled probes tiling the hTR where used (hTR probes: hTR-1/16-R:CCACCCTCCGCAACCC, hTR hTR-43/66-R: GCCCTTCTCAGTTAGGGTTAGAC, hTR-143/166-R: GCTCTAGAATGAACGGTGGAAGG, hTR hTR-304/322-R: CGGCTGA CAGAGCCCAAC, hTR-368/388-R: ACTCGCTCCGTTCCTCTTCC, hTR 407/ 424-R: CGTCCC ACAGCTCAGGG). A single P^32^-labeled MS2 probe was used (MS2 probes: acatgggtgatcctcatgt). The blot was exposed to a PhosphorImager screen and scanned using Image-Quant software.

### TRAP assay

Cell extracts were prepared from cells with CHAPS 1X lysis buffer (10 mM Tris-HCl pH 7.5, 1mM MgCl_2_, 1mM EGTA, 0.1mM Benzamidine, 5mM β mercaptoethanol, 0.5% CHAPS, 10% glycerol). TRAP essay was performed on cells extract with TRAPeze telomerase detection kit (Millipore) according to the manufacturer instruction. The TS primer was radiolabeled with γ P-ATP prior to PCR amplification. DNA amplicons were separated by a 10% poly-acrylamide gel electrophoresis. The gel was dried and exposed with a PhosphorImager screen and scan with Image-Quant.

### Live-cell imaging

For live-cell microscopy, cells were plated on glass-bottom 35mm dishes (Fluorodish FD35-100 WPI) in growth media. Three hours before imaging, growth media was replaced with imaging media (DMEM phenol red free with 10%FBS and 25mM HEPES pH7.4) and cells were imaged on a Zeiss Axio-Observer Z1 Yokogawa CSU-X1 spinning disk inverted confocal microscope equipped with two EMCCD Evolve cameras (Photometrics, 512×512 pixels, 16 µm). The images were acquired with a 100X NA 1.46 oil objective. The pixel size of the images obtained was 0.133µm. Zeiss TempModule was set to 37°C two hours prior to imaging in order to allow all components of the imaging chamber to reach the appropriate temperature. The different lasers (405nm 50mW OPSL, 488nm 100mW diode, and 561nm 40mW diode) were turned on for a minimum of one hour before imaging.

For DualCamera imaging of hTR-GFP and mCherry-hTRF1, cells were excited simultaneously with the 488nm and 561nm lasers. Emission was split by a Zeiss FT 560 beam splitter. The Zeiss DualCamera calibration Wizard software was calibrated with a slide of multi-fluorescent tissue to apply an Affine (translation + rotation + isoscaling) transform to the 561nm laser image. Images were acquired continuously with 70ms exposure time (plus 30 ms transfer, resulting in 100 ms interval) for 400 images. To determine the localization precision of the hTR-GFP and mCherry-hTRF1 imaged in DualCamera mode, 100 nm TetraSpeck beads were suspended in glycerol droplets to achieve diffusion coefficients similar to that of telomeres. The localization precision of the 488nm and 561nm images of the beads was 45 nm.

For short tracks imaging, hTR-GFP was filmed continuously at 100ms exposure time for 100 images on a single Evolve camera (frame rate of 10 images per second). A single image of the mCherry-hTRF1 signal with 350ms exposure was acquired both at the start and the end of the time series. For long tracks, mCherry-hTRF1 and hTR-GFP were acquired sequentially on a single Evolve camera at 5 second intervals for 60 images. GFP was acquired with 100ms exposure time and mCherry with 150ms exposure time. Focus was maintained with the Zeiss Definite Focus LED IR 835nm laser every 10 images.

### Single particle tracking

For single camera images, Zeiss czi files were opened in Image J and saved as single channel 16 bit TIFFs. The exponential fit bleach correction was applied to all image series that showed decay. The outline of the nucleus was drawn with the freehand tool using the mCherry-hCdt1 /mCherry-hTRF1 image and saved as an ROI. Image J TrackMate plugin (Tinevez et al., 2017) was used for single particle tracking within nuclear ROI. The Laplacian of Gaussian spot detection matrix was set to four pixels (0.53 µm) and median filter and sub-pixel localization were selected. TrackMate simple LAP tracker was used with a linking max distance of 0.8 µm, and zero gap-closing. A filter was set such that only tracks with 5 or more consecutive spots were considered. The diffusion coefficient (D) is the slope of the MSD over time. The Mean Square Displacement (MSD) is calculated with the formula:

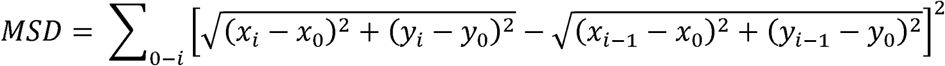

We calculated D as the slope of the MSD over time for the first five points of each track. To calculate D for the thousands of hTR-GFP tracks in the nucleoplasm, an R Studio script was designed to obtain D for all tracks from the TrackMate output.

For Dual Camera images, Zeiss czi files were exported to the OME TIFF format from within the ZEN 3.0 (blue edition) software to conserve the Affine alignment of the second camera images. OME TIFF files were then analyzed in Image J. Exponential fit bleach correction was applied to all image series that showed decay. ImageJ TrackMate plugin was used to track hTR-GFP and mCherry-hTRF1 within a 21 pixel round region of interest (ROI) positioned over a single telomere. The Laplacian of Gaussian spot detection matrix was set to four pixels (0.53 µm) and median filter and sub-pixel localization were selected. TrackMate simple LAP tracker was used with a linking max distance of 0.2 µm, and zero gap-closing.

The formula for the radius of gyration was:

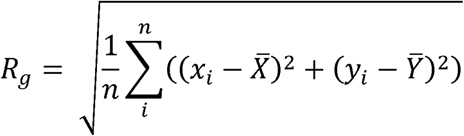

To represent the variation in amplitude movement of hTR and telomeres in DualCamera images, a sliding window of 500 ms moving in increments of 100 ms was applied to either the diffusion coefficient or the radius of gyration, and sliding window values were calculated. The sliding window of diffusion coefficient and radius of gyration of 100 nm TetraSpeck beads in glycerol droplets were calculated to determine the amplitude of variation of perfectly colocalized signals with the DualCamera microscope. The difference between the radius of gyration of the bead in 488nm and 561nm laser images was 0.012µm. For the diffusion coefficient, the average difference between the D in the two colors was 0.01 µm^2^/sec. These values represent the precision of these diffusion measures with the imaging conditions we used.

### Photoactivation

For photoactivation experiments, cells expressing mCherry-hCoilin and MCP-paGFP were plated as described above for live-cell imaging. The cells were imaged with mCherry filter in Live mode. The Cajal body marked with an mCherry-hCoilin was positioned in the center of the field of view (FOV), so that it could be targeted by the centered 1.5 µm diameter illumination area of the Zeiss Direct-FRAP module with the 405nm laser for photoactivation. Five pre-bleach images were acquired. The center of the FOV was then exposed to the 405nm laser for 500ms, and 195 images were then acquired to follow the hTR^5’MS2^-paGFP signal. The time series was filmed continuously at 150ms exposure time (plus 30 ms transfer, resulting in 180ms interval).

Analysis of the photoactivation experiments was carried out by loading Zeiss czi files in Image J and saving then as single channel 16-bit TIFFs. The exponential fit bleach correction was applied to the paGFP image series. A 7 pixels diameter circle (0.93 µm) ROI was used to measure fluorescence grey levels (integrated density) of paGFP in the center of the Cajal body of FOV position. The fluorescence background of paGFP was measured in four 7 pixels diameter circular regions in the nucleoplasm and the average background at each time-point was subtracted from the value of the hTR-paGFP in the CB. The value of fluorescence in the Cajal body center in the first image after the 405nm laser photoactivation, was set to 1. All subsequent values were reported as a fraction of 1. The paGFP fluorescence decay curves were analyzed in GraphPad prism with exponential decay equations. The parental HeLa hTR^WT^ cells had a single population of MCP-paGFP which fits the one phase decay equation with a median half-file of 0.4 seconds.

### Photobleaching

Cells were grown on 35 mm glass bottom (#1.5) dishes (MatTek). Prior to each imaging session, cells were transferred to DMEM lacking phenol red and supplemented with HEPES. Imaging was performed on a Nikon Eclipse Ti2 equipped with a Yokogawa Electric CSU-W1 spinning disk module, and using a 100x objective (NA 1.35, Plan Apo). Image acquisition was performed using Nikon Elements software. A Tokai Hit stage-top incubator was used to control temperature, humidity, and CO2 levels. CBs with colocalizing hTR were identified, and a circle of 2 micron diameter around the Cajal body was bleached with 405 nm beam (Bruker). Five initial images were taken of each cell before bleaching. After bleaching, images were taken every 600 ms with a 100 ms exposure time for 60 s. Images were collected using two Andor iXon 88 Life EMCCD cameras connected by a twin-camera module (Cairn). Cells were illuminated with 488 nm light (for MS2-hTR) and 564 nm light (for mCherry-Coilin). Registration was corrected using Tetra Speck beads (Invitrogen) and images were flat-field corrected for the 488 nm channel using a solution of fluorescein (Model and Burkhardt, 2001). FRAP analysis was performed in FIJI (Schindelin et al., 2012) using the FRAP Profiler plugin (http://worms.zoology.wisc.edu/research/4d/4d.html). FRAP curves were fit to averages of the data sets in Prism using a one-phase association model.

### smiFISH and Super-resolution microscopy

Single molecule RNA FISH against hTR was performed according to the method described in (Tsanov et al., 2016) with some modifications. HeLa cells were plated on acid washed glass coverslips (22×22mm no1.5) and fixed in 4% paraformaldehyde for 20 minutes at room temperature. Cells were then permeabilized in 0.5% Triton-X100 at room temperature for 5 minutes and transferred to 1X SSC 15% formamide solution for hybridization. A 15 hTR smiFISH probe set was designed with Oligostan R script described in (Tsanov et al., 2016) with additional 4 probes within the MS2 sequence. The hTR probe sets were hybridized with a FLAP-Cy5 oligo. Hybridization was carried out overnight at 37°C. Coverslips were washed with 1XSSC;15% formamide then rinsed in PBS and blocked in 0.1% Triton-X100; 2%BSA for 1 hour. Cells were then incubated with coilin antibody (1:500 in blocking solution) for 2 hours at room temperature. Alexa488 anti-mouse secondary antibody (1:6500 in blocking solution) was incubated for 50 minutes at room temperature. Lastly, cells were stained with H33342 (0.5µg/ml in PBS) and mounted with Prolong Glass antifade (ThermoFisher P36982) for 72 hours prior to image acquisition.

hTR smiFISH images were acquired with a 63X NA 1.46 oil objective on a Zeiss Elyra PS.1 super resolution microscope equipped with an Andor EMCCD iXon3 DU-885 CSO VP461 camera (1004×1002 pixels). Lasers used included: 50 mW 405 nm HR diode, 100mW 488 nm HR diode, 150mW 642 nm HR diode were used to image H33342 (DNA stain), AF488 and CY5 dyes respectively. Structured illumination images from z-stack were acquired using three rotations and a grid size of 42 µm for all channels. TetraSpeck microspheres (ThermoFisher T7279 0.1µm) in the same mounting media (Prolong Glass ThermoFisher) as the samples were imaged in all channels to correct for pixel shift. A built-in channel alignment tool in ZEN 2012 SP5 which uses an Affine image alignment algorithm was used for image registration. Maximum intensity projection images are shown in the smiFISH-IF figures, but the hTR spots and CBs were analyzed as volumes with the 3D ImageJ Suite tools(Ollion et al., 2013). The voxel size after the SR-SIM image processing was 0.04µm in X, Y with a step size of 0.18µm in Z. A 3D reconstruction of the nucleus was used to quantify the number of hTR foci. The eccentricity ratio was calculated by measuring the center to center distance between the Cajal body and a hTR foci, divided by the radius of the Cajal body along the same axis.

---

**Figure S1.**
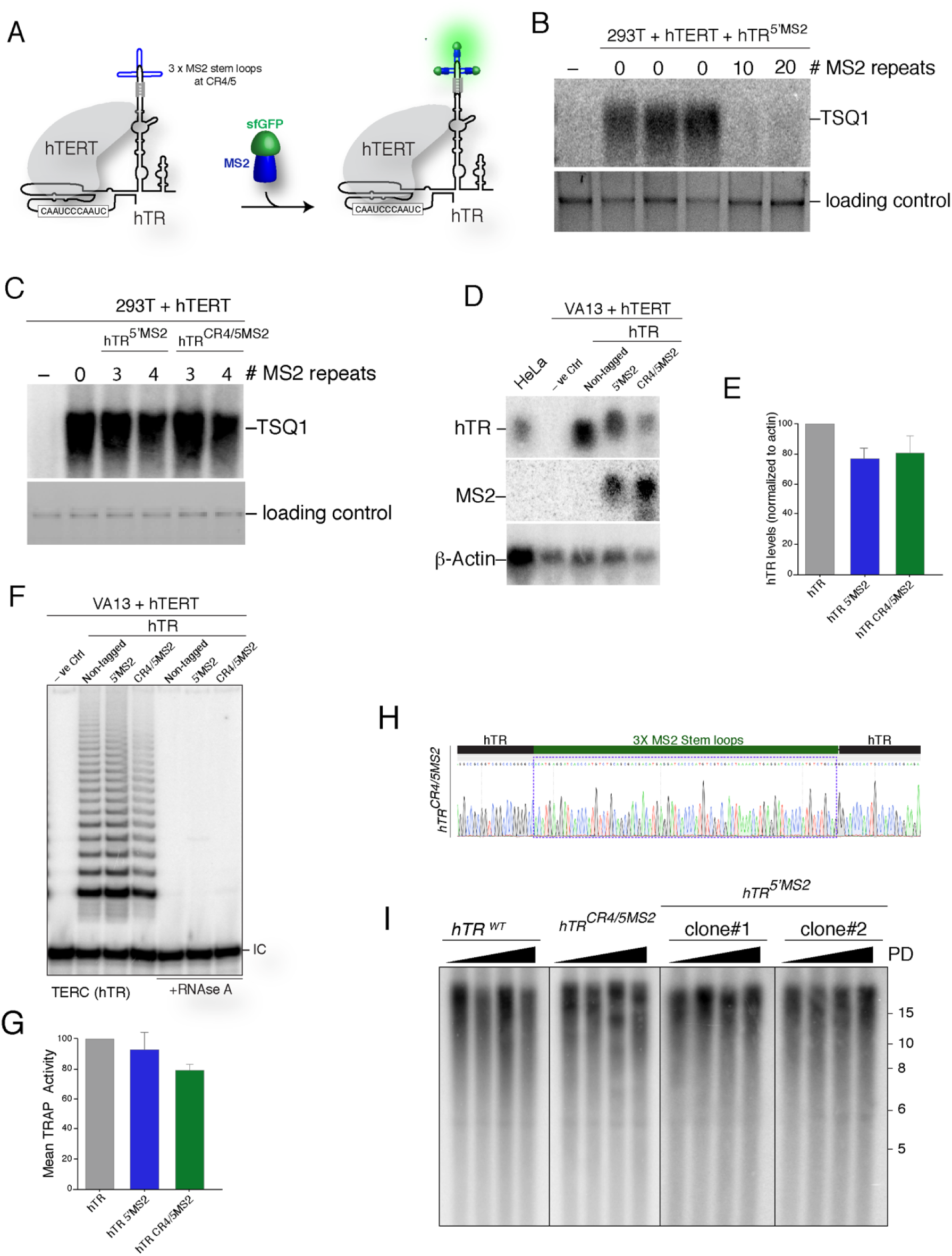
Validation of MS2-tagged hTR, related to Figure 1. **(A)** Schematic depiction of hTR tagging in the CR4/5 region with 3MS2 stem loops that are recognized by GFP-MS2 coat protein. Dual RNA linkers were incorporated to stabilize the folding of the MS2 stem-loops. **(B)** Southern blot analysis detects incorporation of TSQ1 sequences at telomeres in 293T cells overexpressing hTERT and hTR^5’MS2^ with 0, 10, and 20 MS2 stem-loops. **(C)** Detection of TSQ1 sequences at telomeres in 293T cells expressing hTERT and hTR^5’MS2^ or hTR^CR4/5MS2^ with 0, 3, and 4 MS2 stem-loops. **(D)** Northern blot in cells with the indicated treatment. Probes were used to detect MS2, hTR and β-actin (control). **(E)** Quantification of Northern blot in D. **(F)** TRAP assay to detect telomerase activity in VA13 cells with the indicated vectors. **(G)** Quantification of TRAP assay in F. **(H)** Sanger sequencing confirming the integration of 3MS2 stem-loops in the CR4/5 region of the hTR locus. **(I)** Southern blot analysis to monitor telomere length in HeLa 1.3 cells with the indicated genotype for ∼50 populations doublings per genotype.

**Figure S2.**
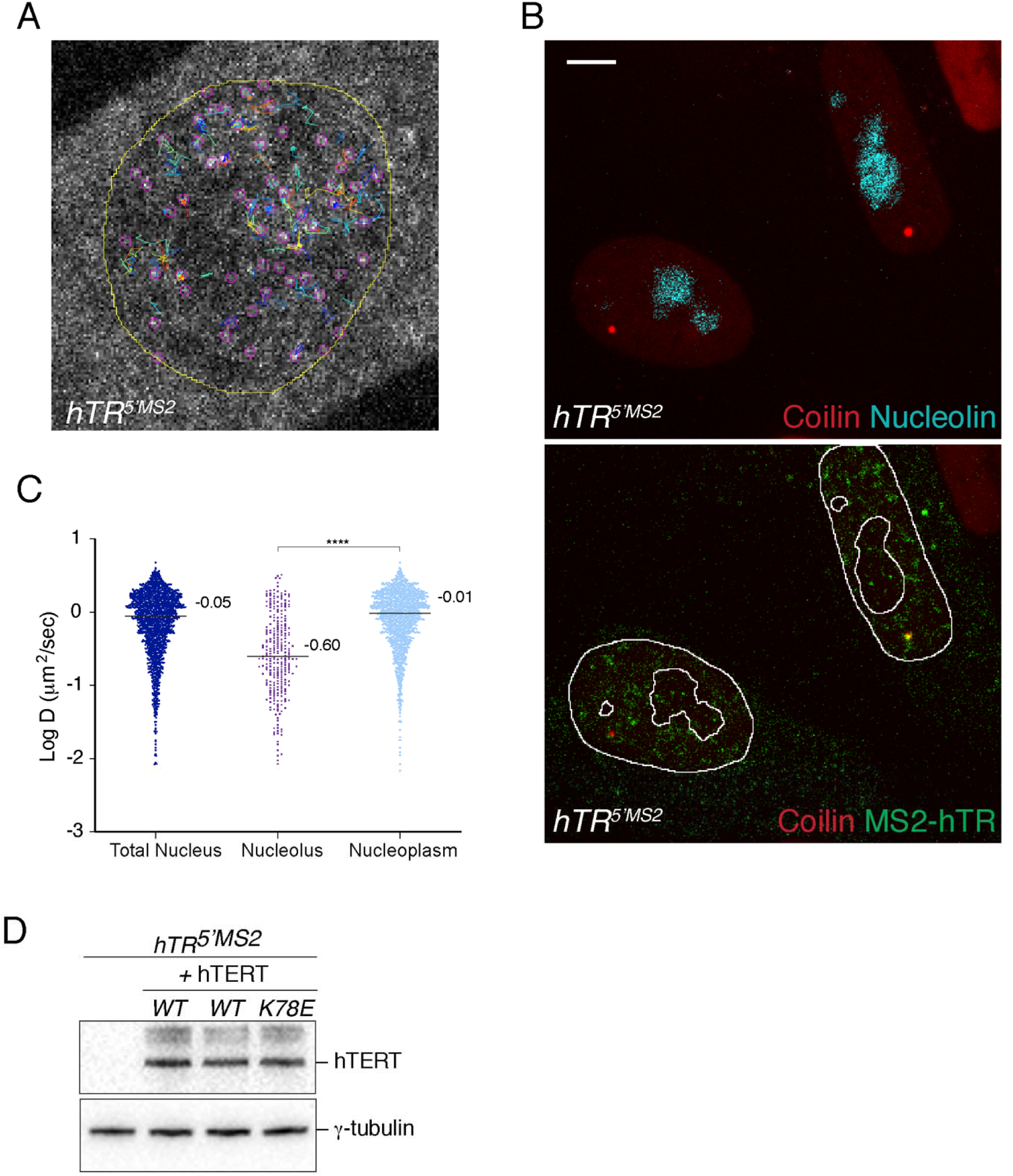
Particle tracking of MS2-hTR, related to Figure 2. **(A)** Examples of MS2-hTR tracks detected by TrackMate. **(B)** hTR particles are detected in the nucleolus. Representative images of MS2-hTR labelled with GFP-MCP (green), nucleolin (blue) and coilin (red). The nuclear membrane, CBs and nucleoli are outlined in white. MS2-hTR particles are visible in the nucleoli. See Movie S4. Scale bar = 5µm. **(C)** Diffusion coefficient of MS2-hTR particles in the total nucleus, nucleolus and nucleoplasm. **** p< 0.001. **(D)** Western blot analysis to monitor expression level of wild-type hTERT and hTERT-K78E in HeLa *hTR^5’MS2^* cells.

**Figure S3.**
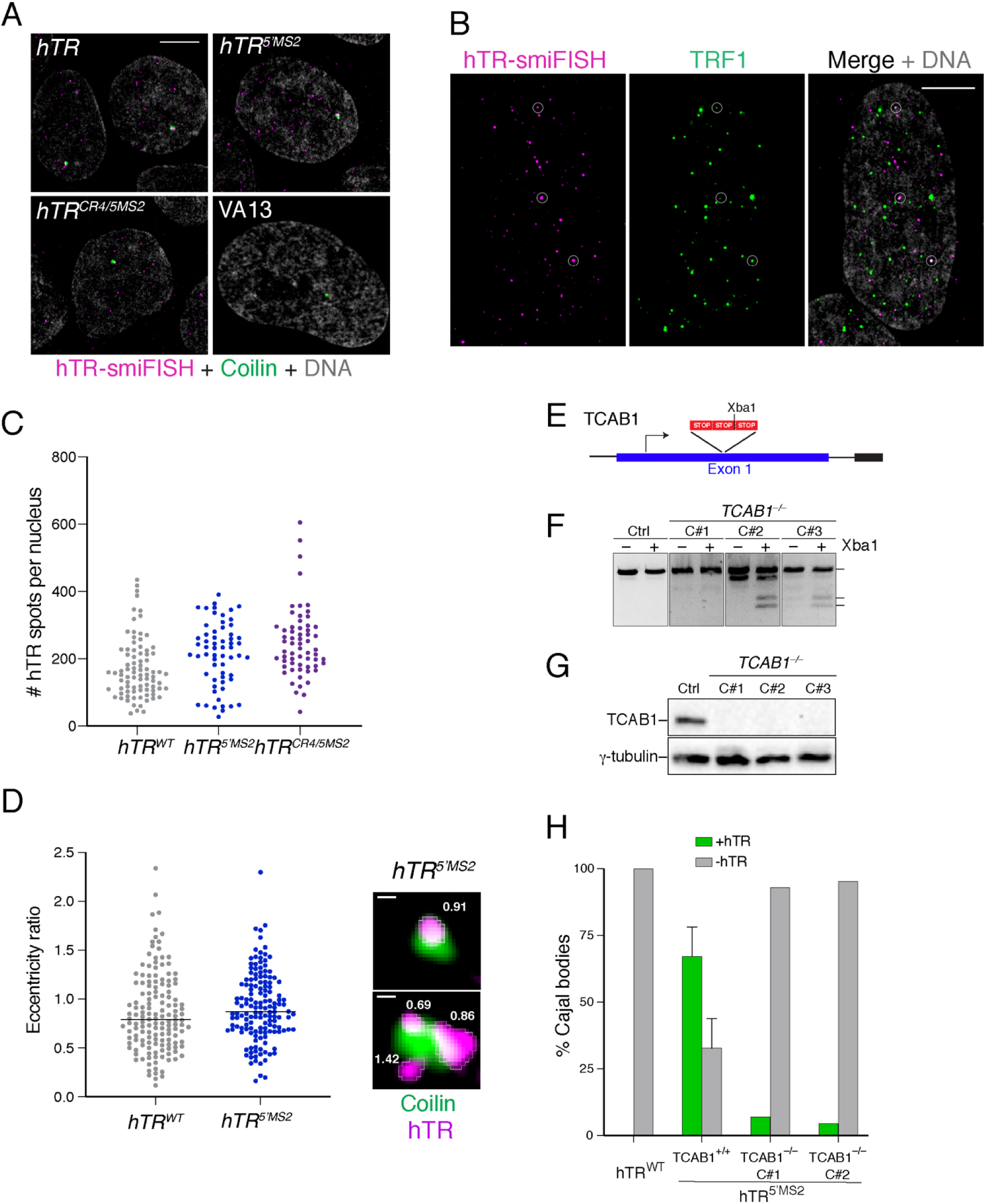
Validation of smiFISH and TCAB1 knockout cell lines for photoactivation experiments, related to Figure 3. **(A)** Representative SIM images of HeLa *hTR^WT^*, *hTR^5’MS2^*, *hTR*^Cr4/5 MS2^ and VA13 cells stained with hTR-smiFISH (15 probes-magenta) and coilin-IF (green). Scale bar: 5 μm. **(B)** Representative SIM images of *hTR^5’MS2^* cells stained with hTR-smiFISH (19 probes-magenta) and mCherry-TRF1 (green). Scale bar: 5 μm **(C)** Quantification of the number of hTR foci per nucleus in *hTR^WT^*, *hTR^5’MS2^* and *hTR^Cr4/5MS2^* cells. **(D)** Quantification of the eccentricity ratio of hTR foci in CBs. Two examples are shown, with their eccentricity ratio indicated. Scale bar = 0.2μm. **(E)** Genome editing strategy to knockout TCAB1 gene in hTR^5’MS2^ HeLa 1.3 cells. A donor cassette coding for three tandem stop codons was introduced by CRISPR/Cas9 gene editing to disrupt the first exon. **(F)** Genotyping PCR on DNA isolated from targeted hTR^5’MS2^ to validate TCAB1 knockout clones. **(G)** Validation of TCAB1 knockout clones by Western blot. **(H)** Quantification of MS2-hTR accumulation in CBs of *hTR^5’MS2^* TCAB1^+/+^ and *hTR^5’MS2^* TCAB1^-/-^ cells using live-cell photoactivation of MCP-paGFP.

**Figure S4.**
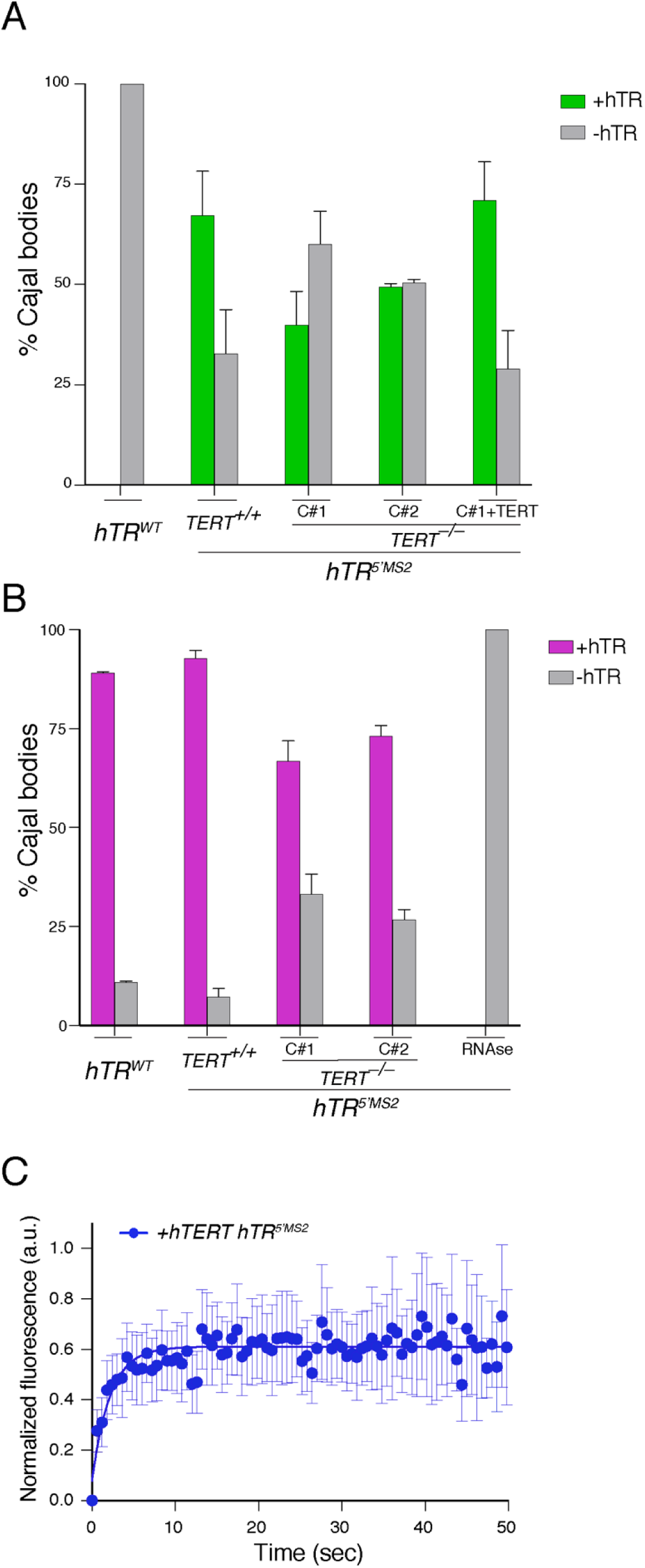
Validation of hTERT knockout clones, related to Figure 4. Quantification of hTR accumulation in CBs of hTR^WT^, hTR^5’MS2^ hTERT^+/+^, two independent clones of hTR^5’MS2^ hTERT^-/-^ cells, and HeLa hTR^5’MS2^ hTERT^-/-^ clone #1 complemented with hTERT using live-cell photoactivation of MCP-paGFP in **(A)** and hTR-smiFISH (19 probes) in **(B)**. **(C)** Fluorescence recovery after photobleaching of MS2-hTR in CBs of *hTR^5’MS2^* cells overexpressing hTERT (N=27 cells).

**Figure S5.**
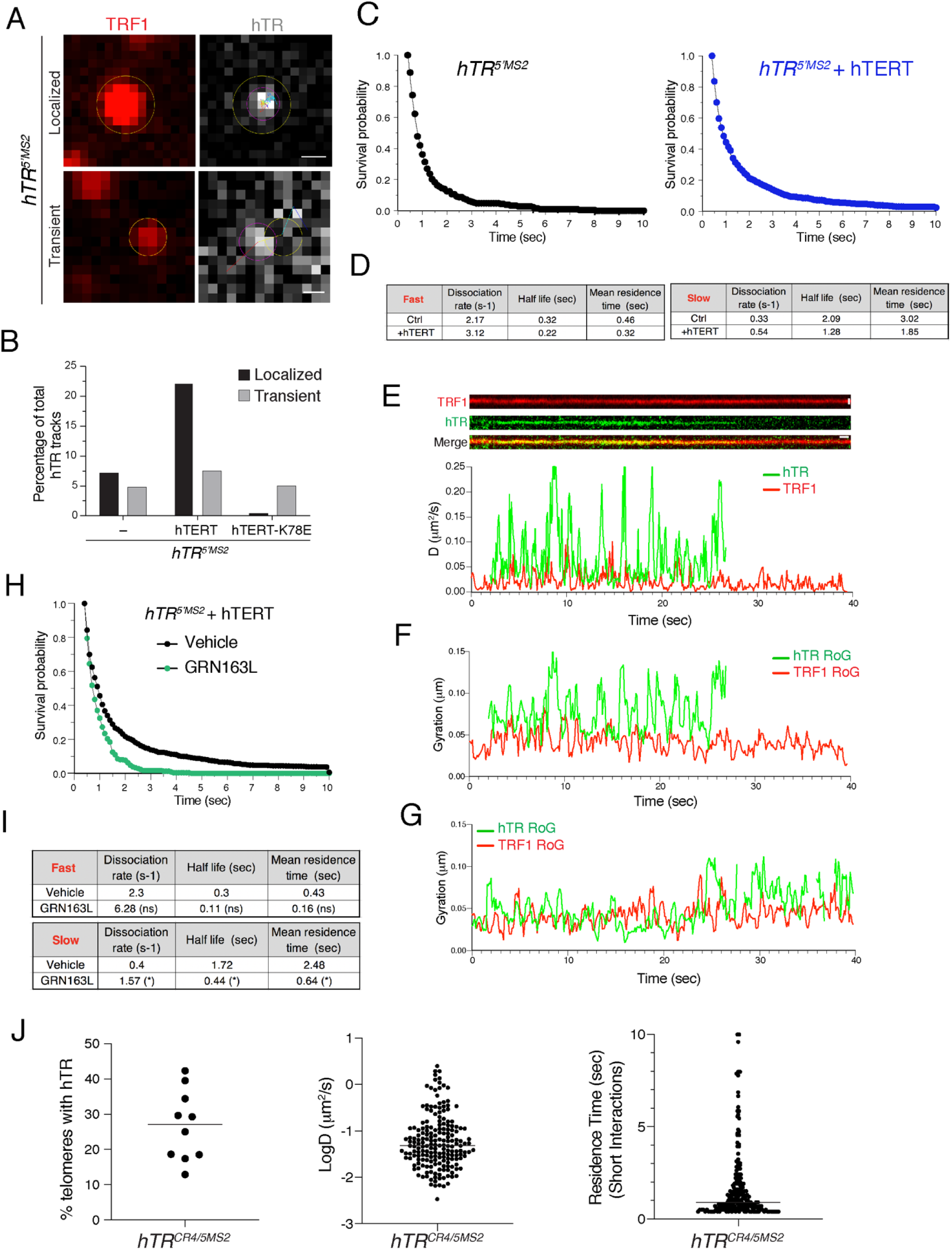
Dual-camera imaging of MS2-hTR and telomeres, Related to Figure 5. **(A)** MS2-hTR particle tracking at telomere (TRF1) reveals localized (top) or transient (bottom) particles at telomeres. Tracks are considered “transient” when an hTR particle remained for only one frame at a telomere, while “localized” tracks correspond to an hTR particle which exhibited telomere interactions that lasted 5 frames or more. Since “transient” hTR tracks were independent of hTERT and persisted in cells expressing hTERT-K78E, they are likely driven by free hTR that is not part of an active telomerase RNP. In contrast, “localized” tracks were significantly increased in the presence of hTERT, but not hTERT-K78E and therefore indicative of productive telomerase-telomere interactions. Yellow circle: telomere position; pink circle: hTR position. Multicolored lines indicate tracks of hTR particles (top scale bar = 0.4µm; bottom scale bar = 0.3 µm). **(B)** Quantification of the percentage of total hTR tracks present at telomeres, either transient or localized, in *hTR^5’MS2^*, *hTR^5’MS2^* + hTERT, and *hTR^5’MS2^* + hTERT K78E cells. **(C)** Survival probability analysis of hTR particles at telomeres in *hTR^5’MS2^* or *hTR^5’MS2^* + hTERT cells fits a bi-exponential distribution. **(D)** Dissociation rate, half-life and mean residence time of fast and slow populations of hTR at telomeres in *hTR^5’MS2^* (Ctrl) or *hTR^5’MS2^* + hTERT cells. **(E)** Top – representative kymograph for hTR and TRF1 tracks. (Vertical scale bar = 0.80 µm, horizontal scale bar = 1 second). Bottom – Running-window analysis of diffusion coefficients at 0.1 second intervals for colocalized hTR (green) and telomere (TRF1, red) over a period of 40 seconds. Notice the loss of MS2-hTR signal at the telomere at t=27 sec. **(F-G),** Running-window analysis of the radius of gyration (RoG) at interval of 0.1 second for colocalized hTR (in green) and telomere (TRF1, red) from tracks depicted in Figure 5F (**F**) or Figure S5E (**G**). **(H)** Survival probability analysis of hTR particles at telomeres in *hTR^5’MS2^* + hTERT cells treated with vehicle (black) or GRN163L (green). **(I)** Dissociation rate, half-life and mean residence time of fast and slow populations of hTR at telomeres in hTR^5’MS2^ + hTERT cells treated with vehicle or GRN163L. (*) p<0.05; (ns) non-significant. **(J)** Validation of MS2-hTR expressed in *hTR^CR4/5MS2^* cells: Left - Percentage of telomeres colocalized with MS2-labelled hTR. Center – Diffusion coefficient of hTR particles at telomeres. Right - Short-term residence time of hTR particles at telomeres.

**Figure S6.**
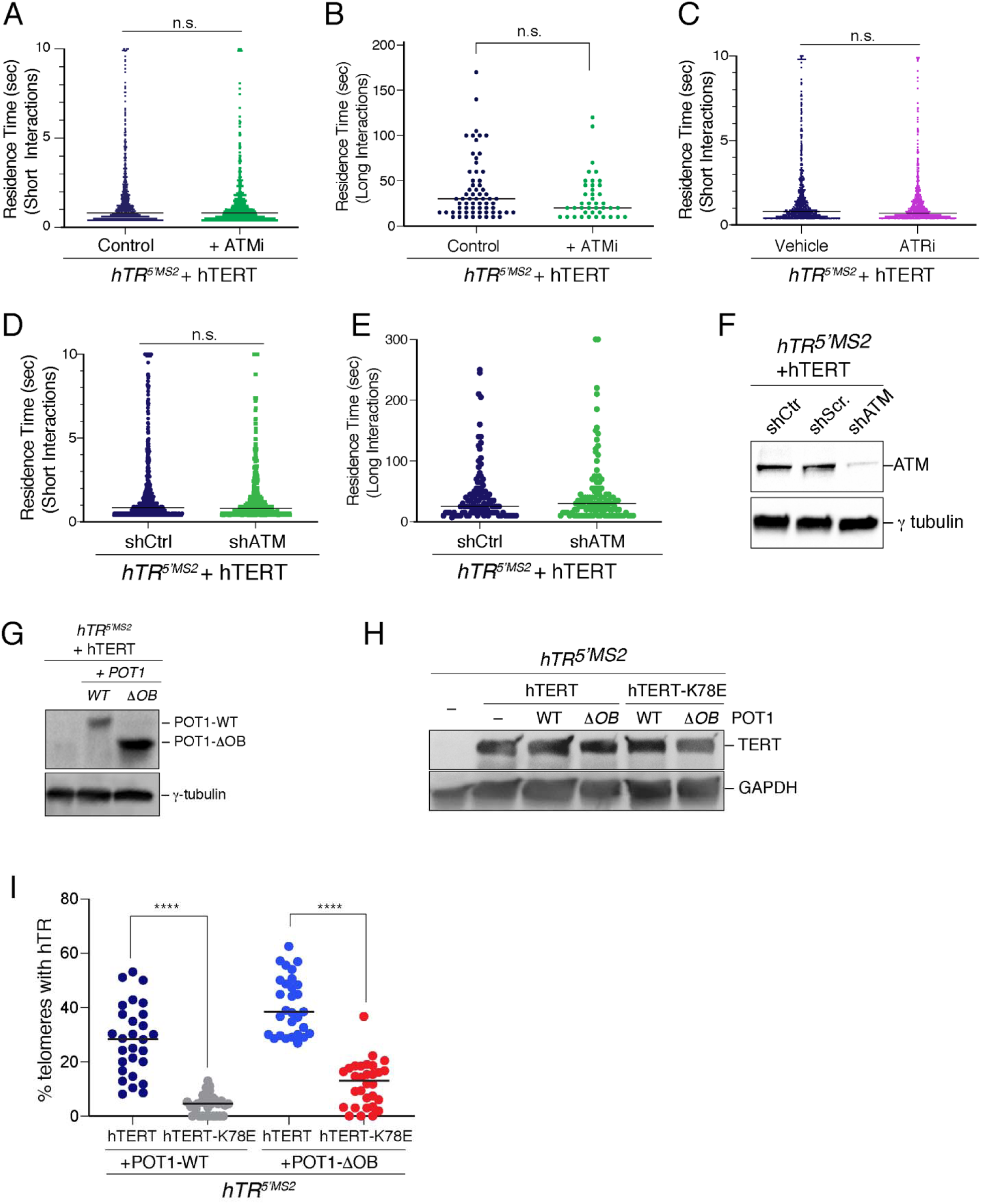
ATM and POT1 regulate telomerase access and residence time at telomeres, Related to Figure 6. **(A)** Short-term residence time of hTR particles at telomeres in *hTR^5’MS2^* cells overexpressing hTERT-WT treated with 3 µM of ATM inhibitor KU60019 (ATMi) or DMSO (control) for 24hrs (N=42 cells for ATMi, N=38 cells for DMSO). **(B)** Long-term residence time of hTR at telomeres in hTR^5’MS2^+hTERT cells treated with 3 µM of KU60019 (ATMi) compared to DMSO (control). **(C)** Short-term residence time of hTR particles at telomeres in *hTR^5’MS2^* cells overexpressing hTERT-WT treated with 0.5 µM of ATR inhibitor VE-822 (ATRi) or DMSO (control) for 3hrs (N=30 cells). **(D)** Short-term residence time of hTR particles at telomeres in *hTR^5’MS2^* cells overexpressing hTERT-WT with either shATM or shCtrl (N=30 cells). **(E)** Long-term residence time of hTR at telomeres in hTR^5’MS2^+hTERT cells with either shATM or shCtrl. **(F)** Western blot analysis of ATM expression in hTR^5’MS2^+hTERT cells expressing either control shRNA (shCtr), scrambled non-targeting shRNA (shScr) or ATM-targeting shRNA (shATM). **(G)** Western blot analysis of POT1-WT and POT1-ΔOB in hTR^5’MS2^+hTERT cells. **(H)** Western blot analysis of hTERT-WT and hTERT-K78E expression in hTR^5’MS2^ cells, hTR^5’MS2^ + POT1-WT cells and hTR^5’MS2^ +POT1-ΔOB cells. (**I**) Percentage of telomeres colocalized with hTR in *hTR^5’MS2^* +hTERT+POT1-WT cells, *hTR^5’MS2^* +hTERT-K78E+POT1-WT cells, *hTR^5’MS2^* +hTERT+POT1-ΔOB cells and *hTR^5’MS2^* +hTERT-K78E+POT1-ΔOB cells. **** p< 0.001.

**Movies S1-S2-S3**

MCP-GFP fluorescence in four cell lines expressing endogenous *hTR^WT^* (control), *hTR^5’MS2^* and *hTR^5’MS2^*+TERT-WT, respectively and co-expressing MCP-GFP. The images were continuously acquired for 10 seconds with an exposure time of 100 msec (scale bar = 5 µm). All the live-cell imaging in the supplementary movies was performed with a spinning disk confocal microscope

**Movie S4**

MCP-GFP fluorescence in *hTR^5’MS2^* cells co-expressing mCherry-Coilin and TagBFP-Nucleolin as shown in supplemental Extended Data Figure 2 panel B. A Cajal body is indicated by a circle, where an accumulation of MS2-hTR is visible. The MCP-GFP images were continuously acquired for 20 seconds with an exposure time of 100 msec (scale bar = 5 µm).

**Movie S5**

Two different movies shown side by side. Photoactivation of MCP-paGFP in CBs of hTR WT cell (left) or hTR^5’MS2^ cell (right). A Cajal body targeted by the 405nm pulse is outlined in white (in red in the first 5 frames). The movies were continuously acquired for 36 seconds with an exposure time of 150 msec (interval 180ms because of 30ms transfer delay). (Scalebar = 10 µm).

**Movie S6**

Representative cell from fluorescence recovery after photobleaching (FRAP) experiments. Dual-camera imaging was used to visualize mCherry-coilin (left) and MCP-GFP bound to hTR (right) simultaneously. hTR foci that colocalize with CBs were identified (arrow), imaged once a second for 5 seconds to establish baseline intensities, bleached, and then allowed to recover while being imaged every 600ms for 1 minute. (Scale bar = 10 µm).

**Movie S7**

Overlay of dual-camera movies. Images of mCherry-hTRF1 (telomeres) and MCP-GFP hTR^5’MS2^ were continuously acquired with dual-cameras for 40 seconds and an exposure time of 70 msec (interval 100ms because of 30ms transfer delay). Arrowhead point to one colocalization event, shown in supplementary Movie S8. (Scale bar = 2 µm).

**Movie S8**

Side by side representation of dual-camera movies. Images of mCherry-hTRF1 (left) and MCP-GFP hTR^5’MS2^ (right) were continuously acquired simultaneously with dual-cameras for 40 seconds and an exposure time of 70 msec (interval 100ms because of 30ms transfer delay). The overlay of tracks was generated with the Trackmate software. (Scale bar = 1µm).

